# End Processing in NHEJ by Polymerase λ and PNKP is coordinated during short-range synapsis

**DOI:** 10.1101/2025.05.25.656038

**Authors:** Alex Vogt, Andrea M. Kaminski, Lars C. Pedersen, Tasmin Naila, Alan E. Tomkinson, Susan P. Lees-Miller, Thomas A. Kunkel, Yuan He

## Abstract

Non-homologous end joining (NHEJ) is a major pathway of DNA double strand break (DSB) repair, capable of directly joining both damaged strands of DNA through the coordinated activities of repair factors that detect the termini, physically bridge them together, and perform the chemistry necessary to complete repair. NHEJ is capable of repairing a variety of damaged DNA, employing various accessory end-processing factors to resolve chemically blocked ends, trim overhangs, and fill gaps in order to achieve directly ligatable DNA ends. To investigate the molecular mechanisms underlying end-processing, we determined the cryo-EM structure of the NHEJ specific polymerase Pol λ bound to the short-range synaptic complex, uncovering the mode of its recruitment to the complex as well as a putative model for its activity. Furthermore, the coordinated end-processing activities of the short-range (SR) synaptic complex simultaneously bound by both Pol λ and PNKP, another accessory factor, demonstrates the ability of NHEJ to form large, multifunctional repair complexes capable of processing a variety of different DNA end structures to effect repair.

## 1 Introduction

Double stranded DNA breaks (DSBs) are the most lethal forms of DNA damage, requiring efficient and accurate repair for cells to retain viability and avoid carcinogenesis^1,2^. Non-homologous end joining (NHEJ) is the major DSB repair pathway in quiescent cells^3^, directly joining both strands of the damaged DNA through the concerted activity of pathway specific factors, that can be roughly categorized into two groups; core NHEJ factors – those required in all instances of repair by NHEJ - and accessory factors, which are required only under certain circumstances^4^. Significant progress has been made in elucidating the molecular mechanisms that underly the activity of the core NHEJ machinery, supporting a multi-step model of repair in which the damaged DNA ends are detected and bound to by the Ku70/80 heterodimer^5^ followed by recruitment of DNA-PKcs^6^, before being brought into a long-range (LR) synaptic configuration mediated by the scaffolding proteins XRCC4, DNA Ligase-IV (LigIV), and XLF^7–9^. Autophosphorylation of DNA-PKcs *in trans* triggers a significant rearrangement of the kinase within the complex precipitating its dissociation^10,11^ and a transition to short-range (SR) synapsis wherein the two copies of LigIV sequentially obtain access to the DSB to perform tandem ligation, resolving the DSB^7,12^.

Due to the variety of lesions produced by different DNA damaging agents, additional chemical processing of the DNA ends is often prerequisite for ligation to proceed^13^. This includes the activities of the NHEJ specific DNA polymerases that fill short gaps and overhangs caused by a loss of nucleotides during damage or earlier stages of repair^14^, nucleases that truncate long overhangs as well as function in special cases of V(D)J recombination mediated by NHEJ^15^, and other DNA repair related enzymes that resolve chemical lesions produced by damage which block direct ligation^16–18^. Single molecule studies in *Xenopus* egg extracts indicate that at least certain end-processing events occur during short-range synapsis^19^, but the molecular details underlying these processes’ coordination with the core NHEJ machinery remain unclear, particularly in terms of the structural accommodation of the accessory factors within the large protein-DNA complexes that mediate repair.

Here we explore the activity of two different end-processing enzymes in the context of the core NHEJ machinery; DNA Polymerase λ (Pol λ), a family X DNA polymerase shown to function in NHEJ by filling gaps and overhangs a few (∼4) nucleotides in length^20^, and Polynucleotide kinase-phosphatase (PNKP), a dual purpose repair enzyme that has kinase activity at 5’-hydroxyl (5’-OH) and phosphatase activity on 3’-phosphate (3’-Pho) DNA termini respectively^16^. We report the cryo-EM structure of the short-range (SR) synaptic complex with two copies of Pol λ bound, elucidating the mechanism of its recruitment to DSB. We also examine the activity of *in vitro* reconstituted complexes to show that both Pol λ and PNKP can cooperatively process the same DNA substrate within a single protein complex. Our results provide evidence for a model in which NHEJ end-processing occurs iteratively during short-range synapsis, with the core NHEJ factors forming a stable platform for the enzymes to flexibly access the DSB junction.

## 2 Results

### Architecture of the Pol λ-NHEJ complex

In order to capture an NHEJ synaptic complex in the presence of Pol λ, we extended the step-wise *in vitro* reconstitution of Ku70/80, XRCC4-LigIV, and XLF^7^ to include both Pol λ and PAXX, a redundant core factor that stabilizes the core NHEJ complexes through interactions with Ku70^9,10^, although its end-bridging activity is not enough to form a synaptic complex capable of ligation in the absence of XLF^10^. PAXX has been shown to physically interact with Pol λ through its globular head domain and stimulate its activity^21^. These factors were all sequentially added to a DNA substrate optimized for complex assembly that contained 1 nucleotide (nt) noncomplementary 3’ overhangs on either side to destabilize binding of LigIV’s catalytic domain and block direct ligation. The sample was then purified and stabilized using GraFix^22^ and subjected to single particle analysis by cryo-EM (methods). From roughly 170,000 particles we obtained a 3D reconstruction with a reported resolution of 7.23 Å (Fig. 1A-B). This relatively modest resolution is in a range similar to that of the previously reported cryo-EM structure of the SR complex (8.4 Å)^7^, and is likely reflective of a degree structural flexibility present in these states. Signal subtraction and local refinement of the XRCC4-LigIV scaffold, distal Ku70/80-Pol λ BRCT, and catalytic site-proximal Ku70/80-Pol λ BRCT respectively yielded reconstructions with nominal resolutions of 4.58 Å, 3.86 Å, and 4.43 Å (Supplementary Fig. 2) respectively allowing us to clearly resolve secondary structure elements of both proteins and DNA.

**Figure 1.**
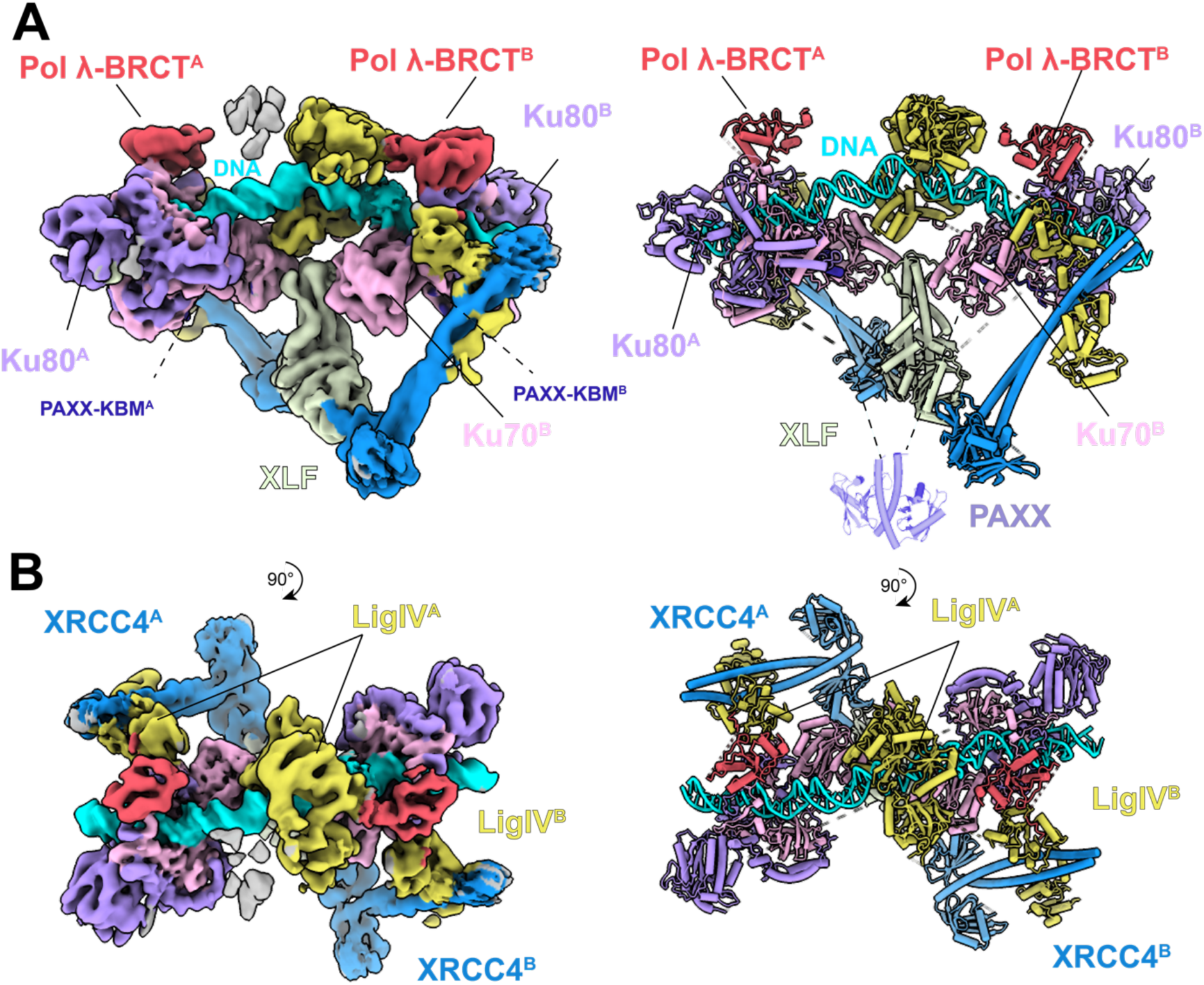
Structure of the SR-Pol λ NHEJ complex. (**A)** Composite cryo-EM map (left) and structural model (right) of the SR-Pol λ complex assembled on a DNA substrate containing symmetrical 1 nucleotide non-complementary 3’ overhangs. (**B)** Top views of the SR-Pol λ complex.

The overall architecture of the SR complex is conserved in the presence of Pol λ, with the XRCC4-XLF-LigIV scaffold and both copies of Ku70/80 adopting almost identical conformations and orientations on the DNA (Fig. 1). Interestingly, more density attributable to XLF’s globular head domains was observed, possibly due to the improvement in map quality in this region compared to previous studies. We were able to model the loop between residues 85-92 (Supplementary Fig. 3A) unreported in the crystal structures, as well as extend the helix that leads into the flexible CTD by 5 residues (Supplementary Fig. 3B). Density corresponding to the DNA is noticeably sharper than reported previously, but the resolution still precludes precisely determining its register. PAXX’s Ku binding motif (residues 179-201) is also present between the core and vWA domains of Ku70 (Supplementary Fig. 3C), as observed in previous studies^9,10^, although its head and stalk domains do not appear in the reconstruction.

### Pol λ is recruited to the NHEJ complex through interactions with its BRCT domain and NTD regions

We observe two copies of Pol λ’s BRCT domain bound to the bridge region of Ku70/80 (Fig. 2A). Upon closer examination, the Pol λ BRCT domain straddles the bridge of Ku70/80 with connective density near residues Arg57 and Leu60, (Fig. 2B) which have been previously shown to be important for Pol λ’s recruitment to DSBs^23^. Given the known dependence of Pol λ’s activity on its BRCT domain^24^, this interaction is essential for anchoring the polymerase to the complex, allowing its catalytic domain more flexible access to the damaged DNA ends (Fig. 2E). Surprisingly, we also see evidence for an interaction between a portion of Pol λ’s N-terminal peptide and the LigIV C-BRCT which bridges XRCC4 and Ku70/80 (Fig. 2C). Guided by AlphaFold3^25^ (AF3) predictions of full-length Pol λ with Ku70/80-LigIV BRCT, we place residues 8-15 at this position. This peptide comprises several highly conserved residues, including Phe10 which interacts with a hydrophobic patch on the BRCT’s surface, and a few positively charged residues that can form an interface with a nearby negatively charged region (Fig. 2D, 2F). This region has also been identified as the polymerase’s nuclear localization signal (NLS)^26^ providing a striking example of the repurposing conserved peptide regions.

**Figure 2.**
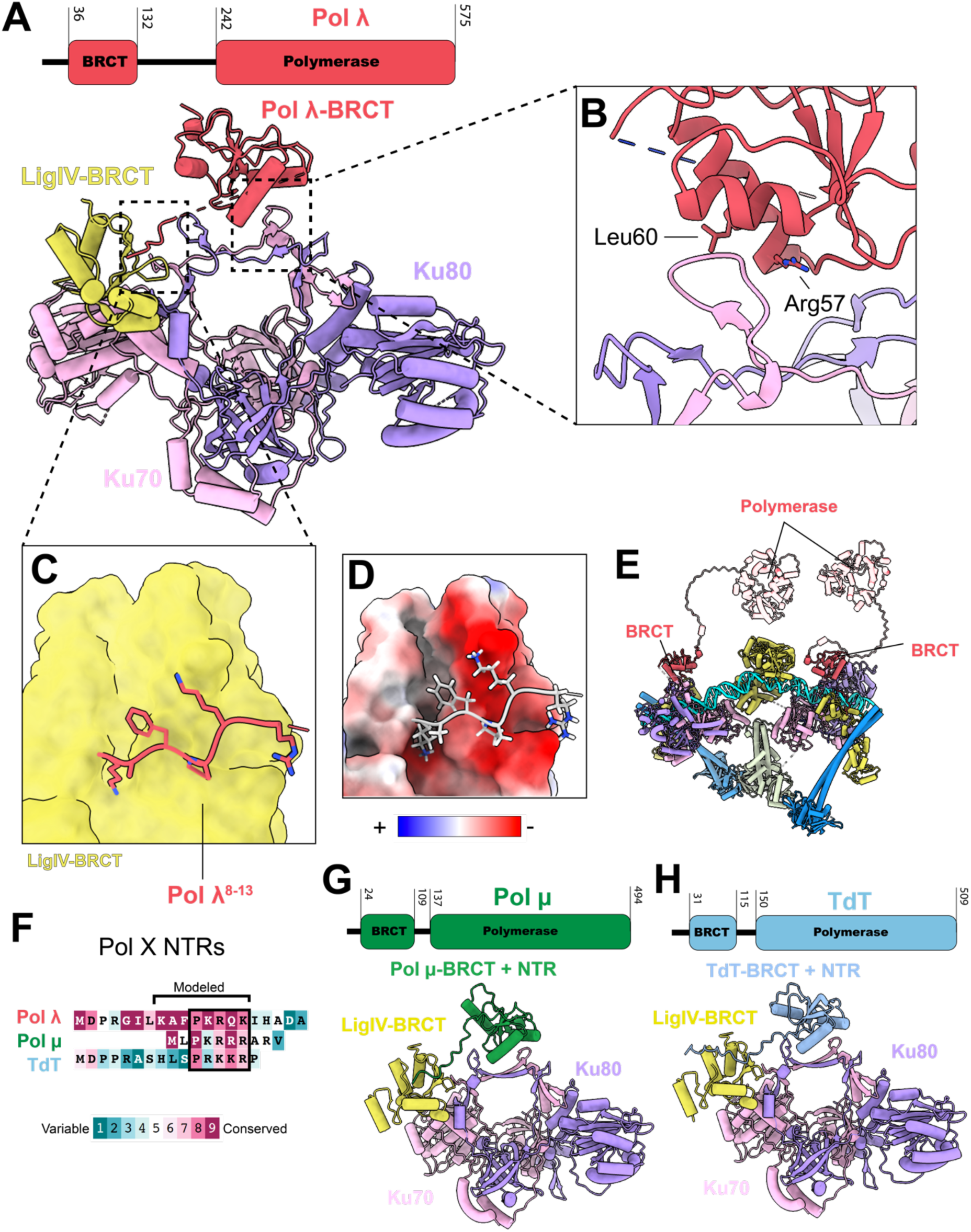
Pol λ recruitment to DSBs and catalytic activity is coordinated with the core NHEJ machinery. **(A)** Model of Pol λ’s NTR and BRCT domain interacting with Ku70/80 and LigIV-BRCT domain. **(B)** Magnified view of the BRCT-Ku70/80 interaction. The position of residues Arg57 and Leu60 are indicated**. (C)** Magnified view of the Pol λ NTR interaction with LigIV-BRCT. **(D)** Magnified view of the Pol λ NTR interaction with LigIV-BRCT colored by electrostatic potential. **(E)** Model showing the flexible tethering of the polymerase domains to the SR complex. Full length AF3 predictions of Pol λ were used to accurately depict the length of the disordered linker region between the BRCT and polymerase domains. **(F)** Sequence alignment of Pol λ, Pol µ, and TDT with a conserved patch of basic residues following a proline indicated. **(G)** Domain architecture and AF3 prediction of Pol µ interacting with Ku70/80 and LigIV-BRCT. **(H)** Domain architecture and AF3 prediction of TdT interacting with Ku70/80 and LigIV-BRCT.

Given the overlapping activities of the other family X polymerases Polymerase µ (Pol µ) and terminal deoxynucleotidyl transferase (TdT), we performed a comparative analysis of AF3 predictions of each with Ku70/80-LigIV BRCT. Both polymerase’s BRCT domains are predicted to bind in a nearly identical manner (Fig. 2G-H) to the bridge of Ku70/80, which is perhaps unsurprising given the structural similarities between these accessory domains. Curiously, regions of both Pol µ and TdT’s N-terminal region (NTR) are predicted to contact LigIV’s BRCT domain as well. Conservation analysis between all three polymerases’ NTRs reveals a patch of conserved positive residues following a conserved proline at residue 11 for both TdT and Pol λ, and residue 3 of Pol µ (Fig. 2F), although residues 8-10 which are represented in our EM map are less well represented. Despite TdT and Pol λ both having longer NTRs than Pol µ, Pol λ’s is much more conserved and predicted to form a more extensive interface with the LigIV-BRCT. It is unclear what role these differences in NTRs play, but given the nearly identical manner in which the BRCT domains appear to bind to Ku70/80, they may help modulate which polymerases are preferentially recruited to the complex through differences in affinity. It is also worth noting the differences in linker length separating the BRCT and catalytic domains of each polymerase, with Pol λ’s being significantly extended (110 residues) compared to Pol µ (18 residues) and TdT (35 residues), which may afford the former much more flexibility in its ability to access the DSB relative to the other family X polymerases.

### LigIV adopts an open conformation to allow end-processing at the DSB

To our surprise, we observed extra density near the bottom of LigIV’s DBD opposite its DNA binding surface. Using AlphaFold3 to screen the factors present in the sample, we were able to identify and model a short portion of the Ku70 CTR at this position (Fig. 3A). Intriguingly, a short helix of the NHEJ specific nuclease Artemis’ CTR binds in this exact position^27^, an interaction which has been shown to play an important role in V(D)J recombination^28^.A comparison of this interaction with the Ku70-LigIV interface reveals a pattern three aromatic residues at conserved positions mediating the interaction (Fig. 3A-B). It is possible that there is some sort of handoff between these factors, perhaps with the Artemis interaction holding LigIV more distant from the DSB during long-range synapsis while Ku70 helps position it at the termini during short-range synapsis.

**Figure 3.**
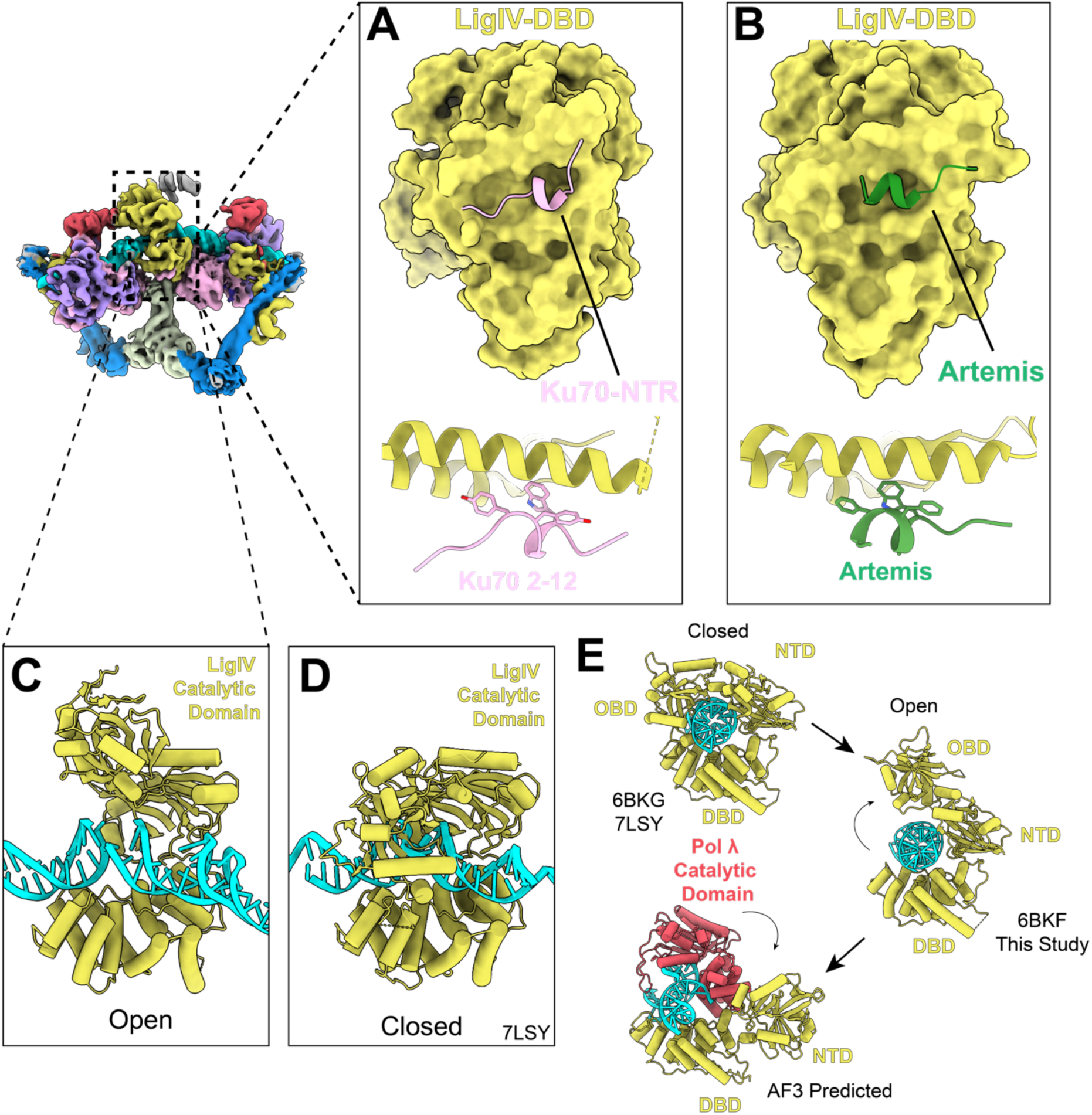
**(A-B)** Comparison of the binding location on the LigIV-DBD of both Ku70 and Artemis peptides with highlighted pattern of three aromatic residues at the LigIV-DBD binding interface shared by Ku70 and Artemis peptides below. **(C-D)** Comparison of the open and closed conformations of LigIV observed between the SR complex captured in this study and previously reported^7^. **(E)** Potential mechanism for Pol λ binding to the DSB based on reported structures of the LigIV catalytic domain and AF3 predictions of the Pol λ-LigIV-DNA complex.

By supplying a DNA substrate with unpaired 1 nt 3’ overhangs, we sought to destabilize the LigIV catalytic domain’s interaction with the DSB. Intriguingly, we still observed the entire LigIV catalytic domain at the DSB, but in an open conformation that aligns well with that reported of LigIV bound to a non-ligatable substrate^29^. This is in contrast to the closed conformation observed in the previously reported SR complex (Fig. 3C-D). In this conformation, the DNA is primarily held by LigIV’s DBD, which serves a dual purpose in the complex by interacting with the vWA domain of the proximal Ku70 to stabilize short-range synapsis, with its NTD forming additional contacts with the DNA. Although the DNA substrate is discontinuous, the double helix is minimally distorted at the DSB junction, suggesting that LigIV is able to accurately align the ends even though they contain noncomplementary bases.

To better understand the mechanism by which Pol λ accesses the DSB junction, we used AF3 to predict potential interactions between the Pol λ catalytic domain, LigIV catalytic domain, and a gapped DNA substrate (Fig. 3E). Based on the available structural information, we hypothesized that the LigIV OBD and NTD domains need to open even further to accommodate Pol λ, potentially being completely displaced by the polymerase’s catalytic domain. This led us to compare predictions on full length LigIV catalytic domain and OBD or OBD+NTD truncations. In line with our hypothesis, the full length LigIV catalytic domain is predicted to assume a slightly more open conformation, while deleting the OBD or OBD and NTD results in a prediction where both Pol λ catalytic domain and LigIV DBD form extensive interactions with the DNA flanking the gap (Fig. 3E). Furthermore, protein-protein interactions are predicted near the helix at 323-328 of Pol λ’s 8 kDA domain. These results support a mechanism whereby the OBD and NTD domains of LigIV’s catalytic domain are displaced by the polymerase during gap filling, while the LigIV DBD maintains its position at the DSB, stabilizing the complex. Also supporting this notion of partial LigIV rather than complete release is the fact that the SR complex does not form when a LigIV construct containing only the BRCT domains is used to assemble the complex, even when the substrates contain terminal complementarity (Supplementary Fig. 4), which indicates Pol λ is alone not enough to stabilize short-range synapsis. Single molecule studies show that mutating or completely deleting the LigIV DBD severely impacts the system’s ability to achieve synapsis^30^, providing further evidence that it may be irreplaceable in this aspect.

### Pol λ allows ligation to proceed on gapped and overhang substrates

To functionally characterize the activity of Pol λ we extended the ligation assay we used previously^7,10^ to include substrates with gaps and overhangs precluding direct ligation. The assembly of the complex and its enzymatic activities are separated by first adding components in a buffer free of Mg^2+^ followed by performing a pulldown on the biotinylated nucleic acid substrate to separate any non-DNA-bound proteins, and resuspending in a buffer containing Mg^2+^ to stimulate ATP hydrolysis, restoring the ligation activity of LigIV (Fig 4a, Supplementary Fig. 5A). Compared to assays where all components are mixed in a bulk reaction, this strategy allows for functional characterization of a more homogenous population of complexes by minimizing the number of freely diffusing factors unbound to DNA, with the added benefit that the sample is amenable for analysis by single particle electron microscopy.

**Figure 4.**
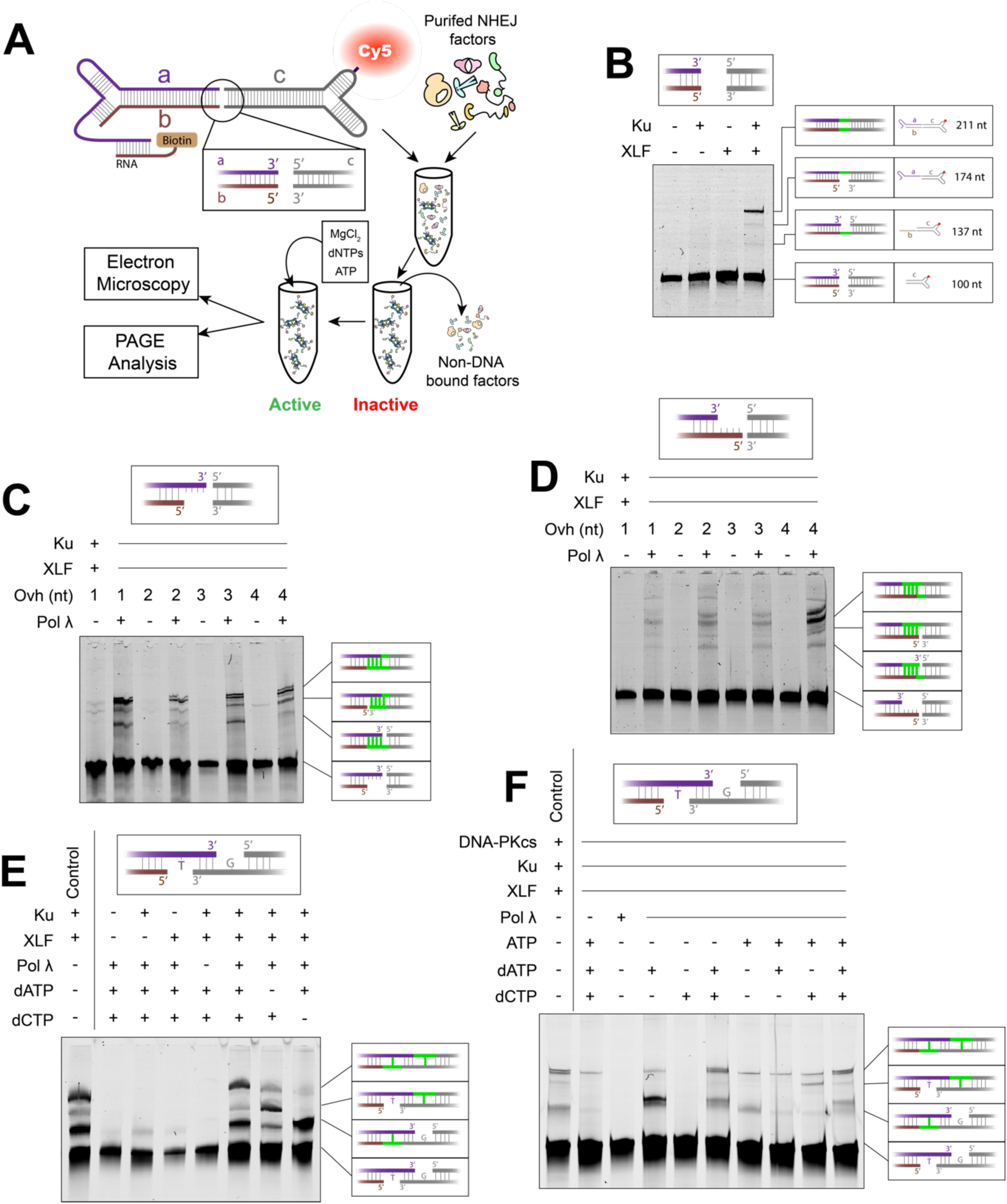
Pol λ facilitates DSB repair on gapped and overhang substrates. (**A**) Schematic showing design of the generic DNA substrate used for complex assembly and protocol for assembly and ligation assay. (**B**) Denaturing TBE/urea gel analysis of end joining of a blunt-end DNA substrate by components of the SR complex. This and all subsequent reactions contain XRCC4-LigIV. Single and double ligation products are indicated on the right with each strand colored the same as in 3A and ligation events colored green. Uncropped gels are available in supplementary material. (**C-D**) Pol λ dependent ligation assay of 1-4 nt 3’ and 5’ overhangs and blunt end DNA. **(E)** Pol λ dependent ligation of a substrate with 3 base-pair microhomology flanked by single nucleotide gaps. Control reaction utilizes directly ligatable substrate with nicks separated by 4 bp terminal complementarity (C4). **(F)** Pol λ dependent ligation in the context of the LR complex demonstrating the use of dATP by DNA-PKcs for phosphorylation (lanes 2 and 4) and the incorporation of ribonucleotides by Pol λ (lanes 7-10). Control substrate is same as in 4E.

On substrates with blunt-ends, all components of the SR complex (Ku70/80, XLF, and XRCC4-LigIV) make up the minimal functional unit capable of ligation (Fig. 4B). When 3’ and 5’ overhangs are introduced on one side, ligation is almost completely ablated, which is recovered by addition of Pol λ and free nucleotides (Fig. 4C-D). Little to no ligation of the strand opposite the overhang is observed in cases where Pol λ is absent, indicating that in this configuration LigIV is inhibited by any missing nucleotides, and Pol λ acts first to produce a blunt end DNA termini prior to ligation. Surprisingly, ligation is also dependent on Pol λ in cases of microhomology flanked by gaps in this system (Fig. 4E), in contrast to previous reports of more flexible LigIV substrate requirements^31^. We note several critical differences in our experimental approach that may account for these discrepancies, primarily the use of a pulldown to purify a relatively homogenous population of stable complexes. In this system, ligation is dependent on the presence of XLF (Fig. 4B), indicative of the formation of the SR complex, which holds the DNA ends in place more rigidly, possibly limiting its ability to accommodate structural distortions caused by ligation over gaps or bubbles. This is in contrast to the flexible synapsis observed in single molecule studies^32^, which is mediated by only Ku and XRCC4-LigIV and may afford more opportunities for LigIV to dynamically interact with the DSB and directly join a wider variety of substrates. In the absence of XLF, this flexibly synaptic state is likely too dynamic to survive pulldown by magnetic beads further illustrating the differences between flexible and short-range synapsis. In our system, as opposed to substrates with no microhomology, single ligation on substrates with terminal base-pairing is possible (Fig. 4E), indicating that in the SR complex LigIV can repair nicks near gaps prior to Pol λ activity, so long as they are flanked by base-paired nucleotides.

Pol λ is also stably recruited to the LR complex (Fig. 4F). As described previously, the two copies of DNA-PKcs present in this configuration protect both DNA ends, and ATP must be supplied to this reaction for the kinases to autophosphorylate *in trans* allowing ligation to proceed (Supplementary Fig. 5B). When assembled on a gapped substrate, ligation by the LR complex is dependent on both ATP and Pol λ (Fig. 4F). Unexpectedly, ligation also proceeds when only dATP is added to a LR-Pol λ complex (Fig. 4F lanes 4 & 6), which indicates that DNA-PKcs can also utilize this cofactor for its kinase activity. Although unusual, there are reports of kinases using dATP as a phosphate donor^33,34^, particularly *in* vitro, although we speculate this is of marginal relevance in the cell where ATP concentrations exceed dATP by orders of magnitude^35^. We also observe Pol λ utilizing ATP for gap filling of a gap with an unpaired thymidine (Fig. 4F lane 7) confirming previous reports of family X polymerases incorporating ribonucleotides during repair^36–38^. We further explored the ability of Pol λ to utilize different dNTPs and rNTPs when filling a single nucleotide overhang, detecting more ligation when the corresponding dNTP was supplied, although products of ligation were observed in the presence of all cofactors (Supplementary Fig. 6). These results are in agreement with previous reports of family X polymerases having medium fidelity, prioritizing restoration of the DNA ends to a directly ligatable configuration over error-free repair.

Structurally, the Pol λ BRCT-Ku70/80 and Pol λ NTR-LigIV BRCT interfaces are not occluded by DNA-PKcs in the LR complex, further indicating that Pol λ’s recruitment to the NHEJ complex is independent of its interaction with DNA and can occur during long-range synapsis. Complementing our findings, a recent study reporting the Pol λ BRCT domain bound to the same position on Ku70/80 NHEJ complexes containing DNA-PKcs^39,40^, providing direct evidence for recruitment of the polymerase at earlier stages of repair. Separating Pol λ’s catalytic domainfrom the BRCT domain by a flexible linker region facilitates the decoupling of recruitment and enzymatic activity, thus allowing the formation of a functional complex without prematurely attempting chemistry at the DSB.

### PNKP is flexibly recruited to the NHEJ complex to repair chemically blocked DNA ends

The end-processing factor PNKP forms a similarly stable complex with the core NHEJ factors (Supplementary Fig. 7A, 7D). However, when subjected to single particle electron microscopy there were no noticeable differences between SR and SR-PNKP complexes in either 2D class averages or 3D reconstructions of negatively stained particles (Supplementary Fig. 7B-C, 7E). It is known that PNKP’s FHA domain interacts with the disordered tail of XRCC4 at the CK2 phosphorylated T233^41,42^, which flexibly tethers it to the complex^43^ precluding structural analysis by electron microscopy.

PNKP’s domain architecture is similar to both LigIV and Pol λ; a flexible linker separates its catalytic domain from an accessory domain that interacts with components of the core NHEJ machinery (Fig. 5A). Therefore, it seems likely that it too is recruited to the complex independently of its direct interaction with the DSB.

**Figure 5.**
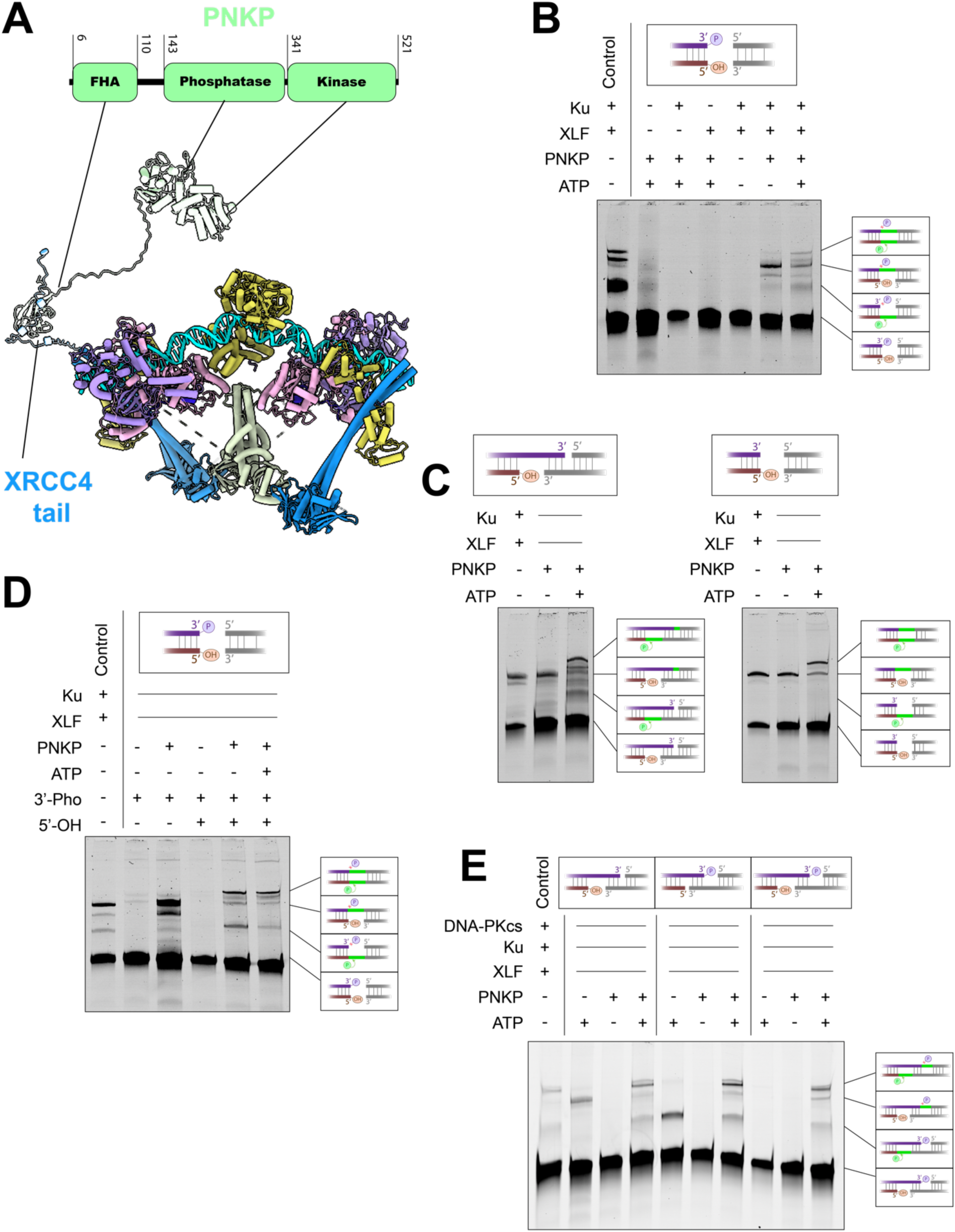
PNKP facilitates repair of chemically blocked DNA termini. **(A)** Domain architecture of PNKP and a putative model for its flexible recruitment to the SR complex via the flexible tail of XRCC4. **(B)** PNKP dependent ligation of a chemically blocked blunt end substrate. End-processing and ligation is dependent on the presence of all components of the SR complex + PNKP. Directly ligatable blunt end substrate is used as control. **(C)** Comparison of ligation on 5’-OH substrates containing with and without microhomology. LigIV is able to carry out single ligation across from the 5’-OH in both cases. **(D)** Comparison of end-processing dependent ligation of 3’-Pho containing substrates. In contrast to those only harboring 5’-OH, the 3’-Pho appears to block functional LigIV engagement with the DSB on substrates without terminal complementarity. Control is same as in 5B. **(E)** PNKP dependent ligation proceeds in an ATP dependent manner in the LR complex. Single ligation proceeds at a nick opposite a 3’-Pho when the lesions are separated by a few base-pairs of complementarity (lane 5). Control substrate is same as used in 4E.

3’-Pho and 5’-OH moieties chemically block ligation by the SR complex that the dual function of PNKP as a DNA phosphatase-kinase is capable of resolving (Fig. 5B). Substrates containing 5’-OH groups, both in blunt end configurations and flanked by microhomology show robust single ligation products at the compatible ends opposite the lesion, demonstrating that the 5’-OH does not inhibit LigIV activity on the opposite strand (Fig. 5C lanes 1,4 Fig. 5E lane 2). However, 3’-Pho severely decreases ligation on the opposite strand (Fig. 5D lane 2), indicating some disruption of the LigIV-DNA interactions necessary to facilitate ligation. The LR complex is also able to recruit PNKP (Fig. 5E), likely because the XRCC4-PNKP-FHA interaction is unperturbed by the presence of DNA-PKcs at the two DNA ends. Like the Pol λ mediated end-processing, addition of ATP is required in all reactions for ligation to proceed in the presence of DNA-PKcs, due to the requirement of its phosphorylation and dissociation from the complex (Fig. 5E).

### End-processing by Pol λ and PNKP is coordinated during short-range synapsis

In cases of DNA damage where both nucleotides are missing and 3’Pho/5’OH groups block ligation the activities of all three enzymes – LigIV, Pol λ, and PNKP – need to be coordinated for repair. Since all three factors form distinct interfaces with core NHEJ factors, there is no structural barrier preventing their recruitment to the same assembly, and we were able to efficiently assemble the entire complex (Supplementary Fig. 8). This SR-Pol λ-PNKP complex processes and ligates substrates containing both overhangs and chemically blocked ends, with double ligation only occurring in the presence of both end-processing factors (Fig. 6A-C). Unexpectedly, when assembled on a substrate containing a 3’-Pho and 4 nt 3’ overhang, Pol λ addition seems to produce substrates that are capable of a single ligation (Fig. 6B-C), although it is unclear how exactly this product is accounted for. In agreement with previous experiments, unpaired nucleotides in the form of overhangs extremely limit ligation, which PNKP alone cannot recover. As expected, the LR complex is also capable of supporting both Pol λ and PNKP (Fig. 6D-E). Introducing microhomology between the gap and 3’-Pho or 5’-OH separates the sites of end-processing, allowing for distinct single ligation products resulting from either Pol λ or PNKP (Fig. 6D lanes 2, 3 Fig. 6E lames 1, 2), while the full range of products are only observed when both factors are present (last lane Fig 6D and 6E).

**Figure 6.**
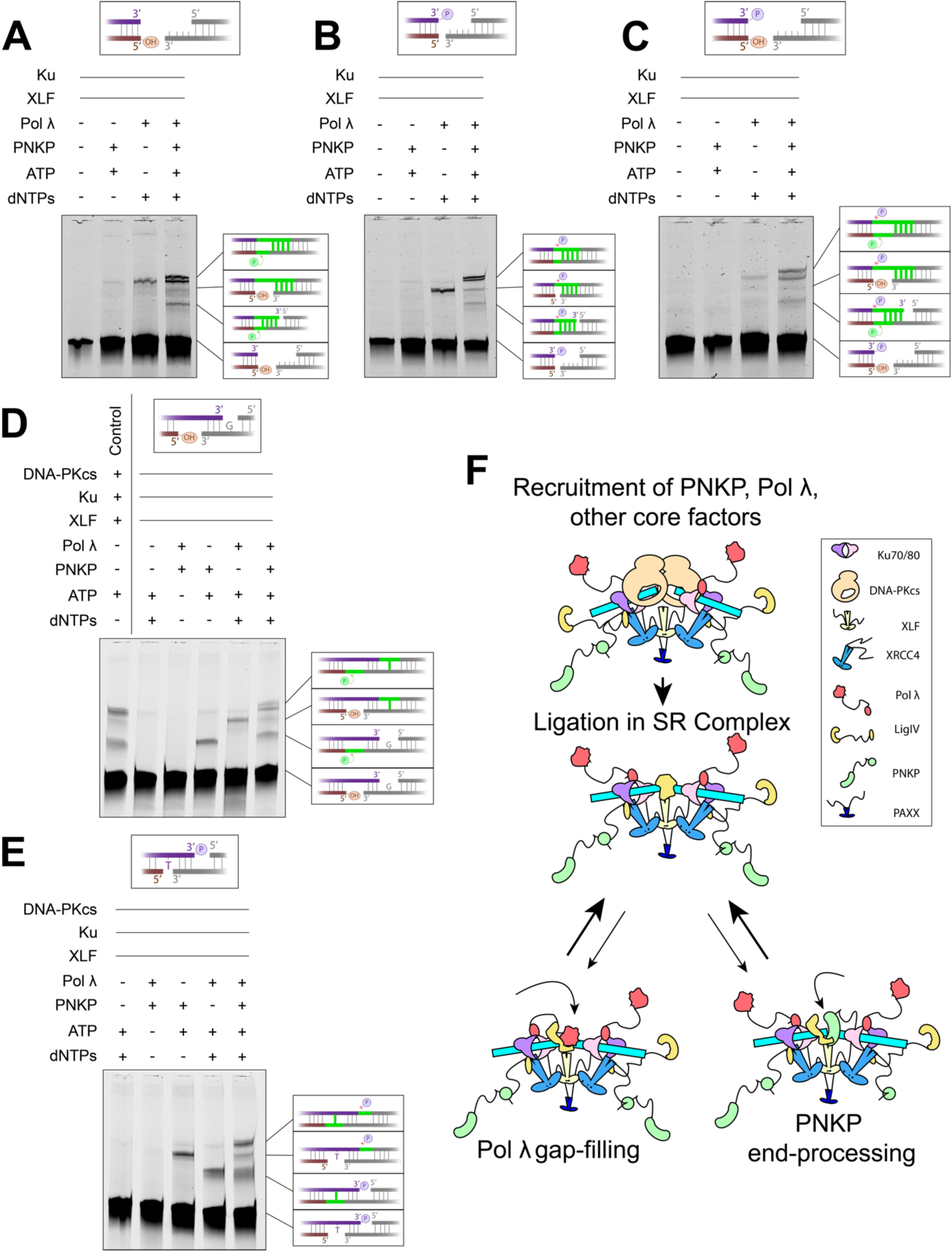
Coordinated DSB processing and repair by Pol, PNKP, and the core NHEJ factors. (A-C) Gap filling and end-processing dependent ligation of incompatible substrates containing 3’ 4 nt overhangs and 3’-Pho and/or 5’-OH. Pol λ appears to have robust activity on substrates with all lesions, although the full range of products are only generated when both factors are present. **(D-E)** Coordinated end-processing dependent ligation of gapped and chemically blocked substrates containing terminal complementarity in the LR complex. Distinct single ligation products dependent on either PNKP or Pol λ are observed (D lanes 2-3, E lanes 1,2), and double ligation products only seen when both factors are present. Control substrate is same as used in Fig. 4E.

## 3 Discussion

The structure of the SR-Pol λ complex and biochemical evidence for the coordinated end-processing by Pol λ and PNKP support a model of NHEJ end-processing where the different factors are recruited at early stages of repair (i.e. pre-synapsis or long-range synapsis) followed by iterative enzymatic activity during short-range synapsis (Fig. 6F). This is consistent with studies showing the importance of the catalytically inert NHEJ polymerases’ BRCT domains and PNKP’s FHA domain in the factors’ ability to resolve DSBs^24,41^. This model is also strongly supported by the recent structural study reporting the Pol λ BRCT binding at the same position of Ku70/80 in the DNA-PK complex^39,40^. All three DNA enzymes described in our investigation (LigIV, Pol λ, and PNKP) contain accessory domains that anchor the protein to the complex and a DNA-binding catalytic domain responsible for performing the actual repair, both of which are separated by flexible linker regions. This domain organization decouples the factors’ recruitment to the complex from the chemistry they perform at the DSB, allowing formation of a multifunctional complex with a variety of enzymes that can dynamically and iteratively access the DSB. Given the important structural role LigIV plays in stabilizing the SR complex, particularly its DBD, it is likely that its disassociation from the DSB is incomplete, consistent with a mechanism of coordinated repair whereby its OBD and NTD domains release the DNA and are displaced by the catalytic domain of another enzyme.

Our results provide mechanistic insight to a growing body of evidence that end-processing, particularly by the NHEJ polymerases and enzymes that resolve chemically blocked ends (PNKP, TDP, etc.), occurs during short-range synapsis in which both ends of the DSB are able to interact with one another. Single molecule studies of this system including both Pol λ and PNKP come to similar conclusions^19^, and crystal structures of Pol λ’s catalytic domains show extensive interactions with both upstream and downstream DNA possible only when both ends of the DSB have come close together^44^. Furthermore, prior to DNA-PKcs eviction, the DNA ends are sterically shielded by DNA-PKcs’ cradle, and access to the termini is occluded by the DBD-helix^7,11^, indicating any enzymatic activity at the DNA ends would have to be well-coordinated with DNA-PKcs. Indeed, such pre-synaptic end-processing by the NHEJ nuclease Artemis has been extensively documented^15,45–47^, but the differences in the properties of this enzyme compared to the other NHEJ end-processing factors (dependence on phosphorylation by DNA-PKcs^48^, extensive interaction surface with DNA-PKcs^49,50^, unique substrate specificities^51^, and central role in V(D)J recombination^52,53^) suggest that its mode of activity may be distinct from other end-processing factors. In contrast, processing by Pol λ and PNKP has no DNA-PKcs requirement and these factors are recruited independently of the kinase, although they are both targets of its phosphorylation^54,55^, the effects of which are currently not well understood. It is possible that limiting these more general end-processing activities to short-range synapsis could enhance fidelity by ensuring the formation of a complete and functional complex before releasing the DNA ends from the protection of DNA-PKcs’ cradle. Occluding access to the damaged termini until they are positioned and primed for close synapsis prevents the filling of any gaps or overhangs until any remaining microhomology is formed, thereby retaining as much original genetic information as possible. The preference for LigIV occupancy at the DSB also favors fidelity by ensuring that ligation is attempted often, resulting in sealing ends as soon as they become compatible, before extraneous end-processing can be attempted.

In summary, we provide structural and biochemical evidence for the coordinated activities of end-processing during NHEJ. Specifically, we identify new interactions between Pol λ and the core NHEJ factors as well as demonstrating the functional cooperation of multiple DNA enzymes on the same DNA substrate in the same complex. By flexibly recruiting different factors to the same complex, NHEJ is able to repair a variety of DSBs while minimizing genetic aberrations. Future studies investigating the activities of other end-processing factors will be needed to determine if this mechanism of coordination is conserved and to test if the formation of even larger complexes capable of repairing a wider variety of DSBs is feasible. Continued dissection of the mechanisms underlying NHEJ’s ability to flexibly repair a variety of DSBs will help further understanding of the phenomena underlying responses to chemotherapy and radiation treatments, with opportunities to develop and optimize targeted therapeutic strategies.

## 4 Methods

### Purification of core and accessory NHEJ protein factors

DNA-PKcs was purified from the nuclear pellet of HeLa cells as previously described^56^ with a measured concentration of 0.5 mg/mL. Previous approaches^57^ were also used to purify Ku70/80, LigIV-XRCC4, and LigIVτιcat-XRCC4 from baculovirus-infected insect cells to final concentrations of 1.53 mg/mL, 7.9 mg/mL and 5.4 mg/mL respectively. Previous methods for XLF^58^, PAXX^57^, and PNKP^54^ expression and purification from *E. coli* were used resulting in concentrations of 2.0 mg/mL, 1.3 mg/mL, and 3.0 mg/mL respectively.

### Cloning, expression, and purification of full-length Pol λ

The sequence encoding full-length human Pol λ (Met1-Trp575) were cloned into the NotI/EcoRI restriction sites of the pGEXM vector^59^ to generate an N-terminal GST-tagged fusion protein. The plasmid was transformed into Rosetta2 (DE3) (Novagen) cells for expression, which was carried out in Terrific broth supplemented with 100 μg/mL ampicillin and 35 μg/mL chloramphenicol. Cultures were grown at 37°C with shaking at 275rpm to an OD_600nm_ of 0.8, when the temperature was reduced to 18°C for 30min. Protein expression was induced by addition of isopropyl-β-D-thiogalactoside (IPTG) to a final concentration of 0.4 mM, and allowed to continue overnight. Bacterial cells were pelleted by centrifugation and lysed by sonication in 25 mM Tris pH 8, 750 mM NaCl, 5% (v/v) glycerol, and 1 mM DTT. The lysate was clarified by centrifugation and bound in-batch to glutathione Sepharose 4B resin (Cytiva). After extensive washing with sonication buffer to remove unbound proteins, the GST tag was removed by on-resin TEV cleavage in 25 mM Tris pH 8, 500 mM NaCl, 5% (v/v) glycerol, 1 mM DTT, rocking gently overnight at 4°C. Cleaved protein was concentrated and further purified by size exclusion chromatography on a Superdex200 (Cytiva) column equilibrated with 25 mM Tris pH 8, 200 mM NaCl, 5% (v/v) glycerol, 1 mM DTT. Fractions containing the purified full-length Pol λ were pooled, concentrated to 4.71 mg/mL in chromatography buffer, and flash frozen in liquid nitrogen.

### Complex assembly and GraFix

All oligonucleotide sequences used in this study are available in table S1. Each was purchased from Integrated DNA Technology (IDT) and annealed as described previously^7^. Substrates were designed to include functional features used in previous investigations^7,10^, and altered primarily at the ends to include overhangs, gaps, 3’-Pho, and 5’-OH to serve as substrates for Pol λ and PNKP.

Complexes were assembled in a buffer containing 50 mM KCl, 10 mM HEPES pH 7.9, 5% glycerol, 1 mM DTT, 100 mM MgCl_2_, and 0.01% NP-40. For negatively stained samples and samples assayed for ligation, approximately 330 nM DNA, 350 nM DNA-PKcs, 430 nM Ku70/80, 1 µM XLF, 800 nM PAXX, 1 µM Pol λ, and 600 nM PNKP were added. Stepwise assembly of the complex was carried by sequentially adding factors followed by 5 min of incubation at room temperature after each step. First Ku was added to each half of the DNA substrate, followed by DNA-PKcs where indicated. Both halves were then mixed together and incubated. Addition of XLF, XRCC4-LigIV, PAXX, Pol λ, and PNKP were all carried out without incubating between factors and the complex was allowed to equilibrate for 10 min at room temperature, followed by addition to a the glycerol-glutaraldehyde gradient or addition of streptavidin-magnetic beads. In cases where GraFix was not used to purify complexes, a streptavidin-based pulldown was performed with 50 nM PAXX included during washing and elution steps where indicated.

GraFix^22^ was used to capture the SR-Pol λ complex. The sample concentration was increased roughly tenfold and the roughly 70 µL sample was then added to the top of a 2 mL 10-30% glycerol/glutaraldehyde gradient, prepared using a BioComp Gradient Master 108. After centrifugation at 30,000 rpm for 16 hrs at 4°C in a TLS-55 rotor using a Optima MAX-TL Ultracentrifuge (Beckman Coulter), the sample was split into 100 µL fractions and the crosslinking reaction was quenched with 500 µM Tris-HCl. Non-denaturing PAGE revealed a peak corresponding to the Pol λ containing NHEJ complex in fractions 12-15, which were subsequently pooled and concentrated to a final volume of 20 µL using Amicron Ultra 0.5 mL centrifugal filters with a 100,000 MW cutoff (Milipore).

### Electron microscopy

Negative staining was carried out as described previously^7^. These samples were imaged using a JEOL 1400 transmission electron microscope equipped with a Gatan 4000 x 4000 charge-coupled device camera operating at 120 kV at a nominal magnification of 30,000x (3.71 Å/pix). Data was collected using Leginon^60^ and SerialEM^61^ with a defocus range of −1.5 µM to −3 µM.

Homemade graphene oxide (GO) grids^62^ were made for cryo-EM by applying GO to glow discharged 2/1, 400 mesh Quantifoil grids that had been glow using a Solarus plasma cleaner (Gatan). Grids were loaded into a Vitrobot IV (FEI) operating at 4°C and 100% humidity, and 4 µL sample was added. Following 5 min incubation on the grid, the sample was blotted with a blot force of −10 for 3 seconds and plunge-frozen in liquid ethane before being transferred to storage in liquid nitrogen.

Two cryo-EM data sets of this sample were collected at the Pacific Northwest Center for cryo-EM (PNCC) totaling 14,845 movies from the same Titan Krios-3 transmission electron microscope (Thermo Fisher Scientific) operating at 300 kV equipped with a K3 direct electron detector (Gatan) and Biocontinuum energy filter with a 10 eV slit width set to 81,000x magnification (0.528 Å/pix in super-resolution mode). 3.162 second, 50 frame movies were collected with a total dose was 62 e^-^/Å^2^ and a defocus range of −2.0 µM to −4.0 µM.

### Image processing and 3D reconstruction

CryoSparc (v4.6)^63^ was used to process both negative stain EM and cryo-EM data. For negative stain, EM images were imported with constant CTF and directly subjected to blob picking, particle extraction, and 2D classification with a box size of 128 pix^2^. To generate the 3D reconstruction of the SR-PNKP complex from negative stain data, multiple iterations of 2D classification yielded 12,199 particles that were used to generate an ab initio model, which underwent homogenous refinement and B-factor sharpening prior to analysis.

For the SR-Pol λ cryo-EM datasets, movies were patch motion corrected and CTF corrected. Initial rounds of blob picking yielded 1,506,486 particles which were extracted with a box size of 360 pix^2^ binned by four and subjected to multiple rounds of 2D classification to generate templates for a second round of picking. Template picking yielded 2,639,430 particles which were extracted with the same box size added to the good particles from the initial round of picking (duplicates were removed). After multiple rounds of 2D classification, exemplary 2D class averages were used to generate low resolution 3D reconstruction ab initio, which was then used as a reference for heterogeneous refinement of all 2,062,285 particles generated from template picking and remaining from blob picking. 4 ‘decoy’ classes generated from discarded particles were also supplied in order to separate junk particles. Iterative 2D classification and heterogeneous refinement using both ‘good’ and ‘junk’ references was performed on particles classified into the particles sorted into the best class, which after plateauing in quality following re-extraction at a box size of 416 pix^2^ retained 77,753 particles resulting in a 8.71 Å reconstruction after non-uniform refinement. Exemplary 2D class averages of this particle set were selected representing common and rare views of the complex, and these were used to train two Topaz^64^ models. Subsequent Topaz extraction yielded 392,215 particles which were combined with the remaining particles from blob and template picking and passed through multiple iterations of 2D classification and heterogeneous refinement. Non-uniform refinement of a final set of 179,057 particles yielded a nominal global resolution of 7.23 Å.

Local refinement was carried out by splitting and masking the complex in three sections; the XLF-XRCC4 scaffold, the distal Ku-Pol λ BRCT-LigIV BRCT domain with the nearby DNA, and the proximal Ku + Pol λ BRCT-LigIV BRCT, LigIV catalytic domain with nearby DNA. UCSF Chimera^65^ and ChimeraX^66^ (made accessible through the SBGrid consortium^67^) were used for volume segmentation and mask generation. All particles were subjected to signal subtraction and locally refined using these masks resulting in nominal resolutions of 4.58 Å for the scaffold, 3.86 Å for the distal Ku-Pol λ, and 4.43 Å for the LigIV catalytic domain + proximal components. Subsequent attempts at 3D classification did not improve the reconstruction.

### Model building

The published cryo-EM model of the SR complex^7^ (7LSY) was used as an initial homology model by rigid body docking into the new cryo-EM map using ChimeraX. The PAXX KBM was placed by aligning the Ku70/80 heterodimer and PAXX KBM portions of the model from the LR-PAXX complex^10^ (8EZA). The AF3^25^ prediction of full-length Pol λ-Ku70/80-LigIV BRCT domains was used to position the Pol λ BRCT domain. AF3 predictions of full-length XLF dimer and Ku70/80-LigIV were positioned in a similar manner, deleting portions where no density was observed. Web 3DNA 2.0^68^ was used to generate a generic DNA model of the 1 nt 3’ overhang substrate used for complex assembly, the termini of which were aligned to the nick in the catalytic site of LigIV. Following rigid body docking of all components, the model was manually rebuilt using ISOLDE^69^ and Phenix real-space refinement^70^.

### *In vitro* ligation assay

Activity assays carried out similar to previously described^7,10^ and overall protocol is diagramed in Fig 4A. Cy5 labeled DNA substrates were annealed and NHEJ complexes were assembled through stepwise addition of the factors shown in a buffer containing 50 mM KCl, 10 mM HEPES pH 7.9, 5% glycerol, 1 mM DTT, and 0.01% NP-40. Samples were then incubated on streptavidin-coated magnetic beads for 15 minutes, following a wash in assembly buffer and resuspension in a reaction buffer identical to the assembly buffer with the addition of 5 mM MgCl_2_. Where indicated, NTPs and dNTPs were added at this step as well a concentration of 1 mM, and the reaction was carried out at room temperature for 15 minutes except where indicated. Samples were then boiled at 95°C for 5 minutes to end the reaction and denature proteins, followed by a 15 minute proteinase K digestion of remaining DNA bound factors for 15 minutes at 37°C. Formamide (50% volume) and 10 mM EDTA were added to samples, which were then boiled for 5 minutes at 95°C before being added to a denaturing urea gel [TBE (tris-borate-EDTA) gel with 8% acrylamide (19:1)]. Gel electrophoresis was performed at 250 V for 30-40 minutes, and gels were scanned using a Sapphire Biomolecular Imager (Azure) at optimal absorbance.

## Acknowledgements

We thank J. Pattie for computer support; J. Meyers, N. Meyer, and M. Miletto at PNCC for data collection and support; M.S.Tsai at the EMB core for assistance in protein expression; and J.P. Lees-Miller for assistance with protein purification. **Funding:** National Institutes of Health grant R01GM135651, and National Institutes of Health grant R01GM144559 to Y.H.; Molecular Biophysics Training Program from NIGMS/NIH (5 T32 GM008382) to A.V.; Northwestern University Structural Biology Facility, NCI CCSG P30 CA060553 (Robert H. Lurie Comprehensive Cancer Center); National Institutes of Health grant P01CA092584 to S.P.L.-M., A.E.T., and Y.H. This work was supported in part by Division of Intramural Research of the National Institute of Environmental Health Sciences, National Institutes of Health [1ZIC ES102645 (to L.C.P.) and Z01 ES065070 (to T.A.K.)]. A portion of this research was supported by NIH grant R24GM154185 and performed at the Pacific Northwest Center for Cryo-EM (PNCC).

## Author contributions

Conceptualization: Y.H., S.P.L-M., A.E.T., and T.K. Methodology: A.V., Y.H., A.M.K., and L.C.P. Investigation: A.V., A.M.K., and T.N. Visualization: A.V. and Y.H. Funding acquisition: Y.H., S.P.L-M., A.E.T., and T.K. Supervision: Y.H. Writing-original draft: A.V. and Y.H. Writing-review and editing: A.V., A.M.K., L.C.P., T.N., S.P.L-M., A.E.T., T.K., and Y.H.

## Data and materials availability

Cryo-EM density maps have been deposited in the Electron Microscopy Data bank (EMDB) under accession numbers EMDB-XXXXX (overall SR-Pol λ complex) EMDB-XXXXX (XLF, XRCCC4, LigIV-BRCT scaffold), EMDB-XXXXX (distal Ku70/80, Pol λ-BRCT), and EMDB-XXXXX (proximal Ku70/80, Pol λ-BRCT, LigiV-catalytic domain). Model coordinates for the SR-Pol λ complex have been deposited in the PDB under accession number XXXX. All data needed to evaluate the conclusions in the paper are present in the paper and/or Supplementary Materials.

## References

1 Stinson, B. M. & Loparo, J. J. Repair of DNA Double-Strand Breaks by the Nonhomologous End Joining Pathway. Annu Rev Biochem 90, 137–164 (2021). 10.1146/annurev-biochem-080320-110356

2 Zhao, B., Rothenberg, E., Ramsden, D. A. & Lieber, M. R. The molecular basis and disease relevance of non-homologous DNA end joining. Nat Rev Mol Cell Biol 21, 765–781 (2020). 10.1038/s41580-020-00297-8

3 Her, J. & Bunting, S. F. How cells ensure correct repair of DNA double-strand breaks. J Biol Chem 293, 10502–10511 (2018). 10.1074/jbc.TM118.000371

4 Ghosh, D. & Raghavan, S. C. Nonhomologous end joining: new accessory factors fine tune the machinery. Trends Genet 37, 582–599 (2021). 10.1016/j.tig.2021.03.001

5 Walker, J. R., Corpina, R. A. & Goldberg, J. Structure of the Ku heterodimer bound to DNA and its implications for double-strand break repair. Nature 412, 607–614 (2001). 10.1038/35088000

6 Sharif, H. et al. Cryo-EM structure of the DNA-PK holoenzyme. Proc Natl Acad Sci U S A 114, 7367–7372 (2017). 10.1073/pnas.1707386114

7 Chen, S. et al. Structural basis of long-range to short-range synaptic transition in NHEJ. Nature 593, 294–298 (2021). 10.1038/s41586-021-03458-7

8 Chaplin, A. K. et al. Cryo-EM of NHEJ supercomplexes provides insights into DNA repair. Mol Cell 81, 3400–3409 e3403 (2021). 10.1016/j.molcel.2021.07.005

9 Seif El Dahan, M., et al. PAXX binding to the NHEJ machinery explains functional redundancy with XLF. Science Advances 9 (2023). 10.1126/sciadv.adg2834

10 Chen, S. et al. Cryo-EM visualization of DNA-PKcs structural intermediates in NHEJ. Science Advances 9 (2023). 10.1126/sciadv.adg2838

11 Chen, X. et al. Structure of an activated DNA-PK and its implications for NHEJ. Mol Cell 81, 801–810 e803 (2021). 10.1016/j.molcel.2020.12.015

12 Graham, T. G., Walter, J. C. & Loparo, J. J. Two-Stage Synapsis of DNA Ends during Non-homologous End Joining. Mol Cell 61, 850–858 (2016). 10.1016/j.molcel.2016.02.010

13 Serrano-Benitez, A., Cortes-Ledesma, F. & Ruiz, J. F. “An End to a Means”: How DNA-End Structure Shapes the Double-Strand Break Repair Process. Front Mol Biosci 6, 153 (2019). 10.3389/fmolb.2019.00153

14 Moon, A. F. et al. The X family portrait: structural insights into biological functions of X family polymerases. DNA Repair (Amst) 6, 1709–1725 (2007). 10.1016/j.dnarep.2007.05.009

15 Ma, Y., Pannicke, U., Schwarz, K. & Lieber, M. R. Hairpin Opening and Overhang Processing by an Artemis/DNA-Dependent Protein Kinase Complex in Nonhomologous End Joining and V(D)J Recombination. Cell 108, 781–794 (2002). 10.1016/S0092-8674(02)00671-2

16 Chappell, C., Hanakahi, L. A., Karimi-Busheri, F., Weinfeld, M. & West, S. C. Involvement of human polynucleotide kinase in double-strand break repair by non-homologous end joining. EMBO J 21, 2827–2832 (2002). 10.1093/emboj/21.11.2827

17 Heo, J. et al. TDP1 promotes assembly of non-homologous end joining protein complexes on DNA. DNA Repair (Amst) 30, 28–37 (2015). 10.1016/j.dnarep.2015.03.003

18 Gomez-Herreros, F. et al. TDP2-dependent non-homologous end-joining protects against topoisomerase II-induced DNA breaks and genome instability in cells and in vivo. PLoS Genet 9, e1003226 (2013). 10.1371/journal.pgen.1003226

19 Stinson, B. M., Moreno, A. T., Walter, J. C. & Loparo, J. J. A Mechanism to Minimize Errors during Non-homologous End Joining. Mol Cell 77, 1080–1091 e1088 (2020). 10.1016/j.molcel.2019.11.018

20 Garcia-Diaz, M. et al. DNA Polymerase, a Novel DNA Repair Enzyme in Human Cells. Journal of Biological Chemistry 277 (2002). 10.1074/jbc.M111601200

21 Craxton, A. et al. PAXX and its paralogs synergistically direct DNA polymerase λ activity in DNA repair. Nat Commun 9 (2018). 10.1038/s41467-018-06127-y

22 Stark, H. GraFix: Stabilization of Fragile Macromolecular Complexes for Single Particle Cryo-EM. Methods in Enzymology 481, 109–126 (2010).

23 Mueller, G. A. et al. A comparison of BRCT domains involved in nonhomologous end-joining: introducing the solution structure of the BRCT domain of polymerase lambda. DNA Repair (Amst) 7, 1340–1351 (2008). 10.1016/j.dnarep.2008.04.018

24 Ma, Y. et al. A biochemically defined system for mammalian nonhomologous DNA end joining. Mol Cell 16, 701–713 (2004). 10.1016/j.molcel.2004.11.017

25 Abramson, J. et al. Accurate structure prediction of biomolecular interactions with AlphaFold 3. Nature 630, 493–500 (2024). 10.1038/s41586-024-07487-w

26 Kirby, T. W., Pedersen, L. C., Gabel, S. A., Gassman, N. R. & London, R. E. Variations in nuclear localization strategies among pol X family enzymes. Traffic 19, 723–735 (2018). 10.1111/tra.12600

27 Ochi, T., Gu, X. & Blundell, T. L. Structure of the Catalytic Region of DNA Ligase IV in Complex with an Artemis Fragment Sheds Light on Double-Strand Break Repair. Structure 21, 672–679 (2013). 10.1016/j.str.2013.02.014

28 Malu, S. et al. Artemis C-terminal region facilitates V(D)J recombination through its interactions with DNA Ligase IV and DNA-PKcs. J Exp Med 209, 955–963 (2012). 10.1084/jem.20111437

29 Kaminski, A. M. et al. Structures of DNA-bound human ligase IV catalytic core reveal insights into substrate binding and catalysis. Nat Commun 9, 2642 (2018). 10.1038/s41467-018-05024-8

30 Stinson, B. M. C., S. M.; Walter, J. C,; Loparo, J. J. Structural role for DNA Ligase IV in promoting the fidelity of non-homologous end joining. Nat Commun 15 (2024). 10.1038/s41467-024-45553-z

31 Gu, J. et al. XRCC4:DNA ligase IV can ligate incompatible DNA ends and can ligate across gaps. EMBO J 26, 1010–1023 (2007). 10.1038/sj.emboj.7601559

32 Zhao, B. et al. The essential elements for the noncovalent association of two DNA ends during NHEJ synapsis. Nat Commun 10, 3588 (2019). 10.1038/s41467-019-11507-z

33 Payne, J. M. & Dahmus, M. E. Partial Purification and Characterization ofTwo Distinct Protein Kinases ThatDifferentially Phosphorylate the Carboxyl-terminal Domain of RNA Polymerase SubunitIIa. J Biol Chem 268, 80–87 (1992). 10.1016/S0021-9258(18)54117-X

34 Das, U., Wang, L. K., Smith, P. & Shuman, S. Structural and Biochemical Analysis of the Phosphate Donor Specificity of the Polynucleotide Kinase Component of the Bacterial Pnkp•Hen1 RNA Repair System. Biochemistry 52, 4734–4743 (2013). 10.1021/bi400412x

35 Traut, T. W. Physiological concentrations of purines and pyrimidines. Molecular and Cellular Biochemistr 140, 1–22 (1994). 10.1007/BF00928361

36 Moon, A. F. et al. Structural accommodation of ribonucleotide incorporation by the DNA repair enzyme polymerase Mu. Nucleic Acids Res 45, 9138–9148 (2017). 10.1093/nar/gkx527

37 Gosavi, R. A., Moon, A. F., Kunkel, T. A., Pedersen, L. C. & Bebenek, K. The catalytic cycle for ribonucleotide incorporation by human DNA Pol λ. Nucleic Acids Res 40, 7518–7527 (2012). 10.1093/nar/gks413

38 Pryor, J. M. et al. Ribonucleotide incorporation enables repair of chromosome breaks by nonhomologous end joining. Science 361, 1126–1129 (2018). 10.1126/science.aat2477

39 Frit, P. et al. DNA polymerase Lambda is anchored within the NHEJ synaptic complex1 via Ku70/80. bioRxiv (2024). 10.1101/2024.08.12.607588

40 Frit, P. et al. Structural and functional insights into the interaction between Ku70/80 and Pol X family polymerases in NHEJ. Nat Commun 16 (2025). 10.1038/s41467-025-59133-2

41 Koch, C. A. et al. Xrcc4 physically links DNA end processing by polynucleotide kinase to DNA ligation by DNA ligase IV. EMBO J 23, 3874–3885 (2004). 10.1038/sj.emboj.7600375

42 Mani, R. S. et al. Dual Modes of Interaction between XRCC4 and Polynucleotide Kinase/Phosphatase. J Biol Chem 285, 37619 –33762 (2010). 10.1074/jbc.M109.058719

43 Aceytuno, R. D. et al. Structural and functional characterization of the PNKP-XRCC4-LigIV DNA repair complex. Nucleic Acids Res 45, 6238–6251 (2017). 10.1093/nar/gkx275

44 Garcia-Diaz, M. et al. A Structural Solution for the DNA Polymerase lambda-Dependent Repair of DNA Gaps with Minimal Homology. Mol Cell 13, 561–572 (2004). 10.1038/s41467-020-18506-5

45 Ma, Y. et al. The DNA-dependent Protein Kinase Catalytic Subunit Phosphorylation Sites in Human Artemis. J Biol Chem 280, 33839 –33846 (2005). 10.1074/jbc.M507113200

46 Chang, H. H., Watanabe, G. & Lieber, M. R. Unifying the DNA end-processing roles of the artemis nuclease: Ku-dependent artemis resection at blunt DNA ends. J Biol Chem 290, 24036–24050 (2015). 10.1074/jbc.M115.680900

47 Gerodimos, C. A., Chang, H. H. Y., Watanabe, G. & Lieber, M. R. Effects of DNA end configuration on XRCC4-DNA ligase IV and its stimulation of Artemis activity. J Biol Chem 292, 13914 –13924 (2017). 10.1074/jbc.M117.798850

48 Niewolik, D. & Schwarz, K. Physical ARTEMIS:DNA-PKcs interaction is necessary for V(D)J recombination. Nucleic Acids Res 50, 2096–2110 (2022). 10.1093/nar/gkac071

49 Liu, L. et al. Autophosphorylation transforms DNA-PK from protecting to processing DNA ends. Mol Cell 82, 177–189 e174 (2022). 10.1016/j.molcel.2021.11.025

50 Watanabe, G., Lieber, M. R. & Williams, D. R. Structural analysis of the basal state of the Artemis:DNA-PKcs complex. Nucleic Acids Res 50, 7697–7720 (2022). 10.1093/nar/gkac564

51 Chang, H. H. Y. & Lieber, M. R. Structure-Specific nuclease activities of Artemis and the Artemis: DNA-PKcs complex. Nucleic Acids Res 44, 4991–4997 (2016). 10.1093/nar/gkw456

52 Moshous, D. et al. Artemis, a Novel DNA Double-Strand Break Repair/V(D)J Recombination Protein, Is Mutated in Human Severe Combined Immune Deficiency. Cell 105, 177–186 (2001). 10.1016/S0092-8674(01)00309-9

53 Mansilla-Soto, J. & Cortes, P. VDJ Recombination: Artemis and Its In Vivo Role in Hairpin Opening. J Exp Med 197, 543–547 (2003). 10.1084/jem.20022210

54 Zolner, A. E. et al. Phosphorylation of polynucleotide kinase/ phosphatase by DNA-dependent protein kinase and ataxia-telangiectasia mutated regulates its association with sites of DNA damage. Nucleic Acids Res 39, 9224–9237 (2011). 10.1093/nar/gkr647

55 Sastre-Moreno, G. et al. Regulation of human pollambda by ATM-mediated phosphorylation during non-homologous end joining. DNA Repair (Amst) 51, 31–45 (2017). 10.1016/j.dnarep.2017.01.004

56 Goodarzi, A. A. & Lees-Miller, S. P. Biochemical characterization of the ataxia-telangiectasia mutated (ATM) protein from human cells. DNA Repair 3, 753–767 10.1016/j.dnarep.2004.03.041

57 Hammel, M. et al. An Intrinsically Disordered APLF Links Ku, DNA-PKcs, and XRCC4-DNA Ligase IV in an Extended Flexible Non-homologous End Joining Complex. *J Biol Chem* **291**, 26987-27006 (2016). 10.1074/jbc.M116.751867

58 Yu, Y. et al. DNA-PK and ATM phosphorylation sites in XLF/Cernunnos are not required for repair of DNA double strand breaks DNA Repair 7, 1680–1692 (2008). 10.1016/j.dnarep.2008.06.015

59 Moon, A. F. et al. Sustained active site rigidity during synthesis by human DNA polymerase μ. Nat. Struct. Mol. Biol. 21, 253–260 (2014). doi:10.1038/nsmb.2766

60 Suloway, C. et al. Automated molecular microscopy: The new Leginon system. J Struct Bio 151, 41–60 (2005). 10.1016/j.jsb.2005.03.010

61 Mastronarde, D. N. Automated electron microscope tomography using robust prediction of specimen movements. J Struct Bio 152, 36–51 (2005). 10.1016/j.jsb.2005.07.007

62 Patel, A., Toso, D., Litvak, A. & Nogales, E. Efficient graphene oxide coating improves cryo-EM sample preparation and data collection from tilted grids. bioRxiv (2021). 10.1101/2021.03.08.434344

63 Punjani, A., Rubinstein, J. L., Fleet, D. J. & Brubaker, M. A. cryosPArc: algorithms for rapid unsupervised cryo-em structure determination. Nat Meth 14, 290–296 (2017). 10.1038/nmeth.4169

64 Bepler, T. et al. Positive-unlabeled convolutional neural networks for particle picking in cryo-electron micrographs. Nat Meth 16, pages 1153–1160 (2019). 10.1038/s41592-019-0575-8

65 Pettersen, E. F. et al. UCSF Chimera—A visualization system for exploratory research and analysis. J Comp Chem 25, 1605–1612 (2004). 10.1002/jcc.20084

66 Pettersen, E. F. et al. UCSF ChimeraX: Structure visualization for researchers, educators, and developers. Protein Sci 30, 70–82 (2020). 10.1002/pro.3943

67 Morin, A. et al. Cutting Edge: Collaboration gets the most out of software. eLife (2013). 10.7554/eLife.01456

68 Li, S., Olson, W. K. & Lu, X.-J. Web 3DNA 2.0 for the analysis, visualization, and modeling of 3D nucleic acid structures. Nucleic Acids Res 41, W26–W34 (2019). 10.1093/nar/gkz394

69 Croll, T. I. ISOLDE: a physically realistic environment for model building into low-resolution electron-density maps Acta Crystallogr D Struct Biol 74, 519–530 (2018). 10.1107/S2059798318002425

70 Afonine, P. V. et al. Real-space refinement in PHENIX for cryo-EM and crystallography. Acta Crystallogr D Struct Biol 74, 531–544 (2018). 10.1107/S2059798318006551

