## Supplementary Information for "End Processing in NHEJ by Polymerase λ and PNKP is coordinated during short-range synapsis"

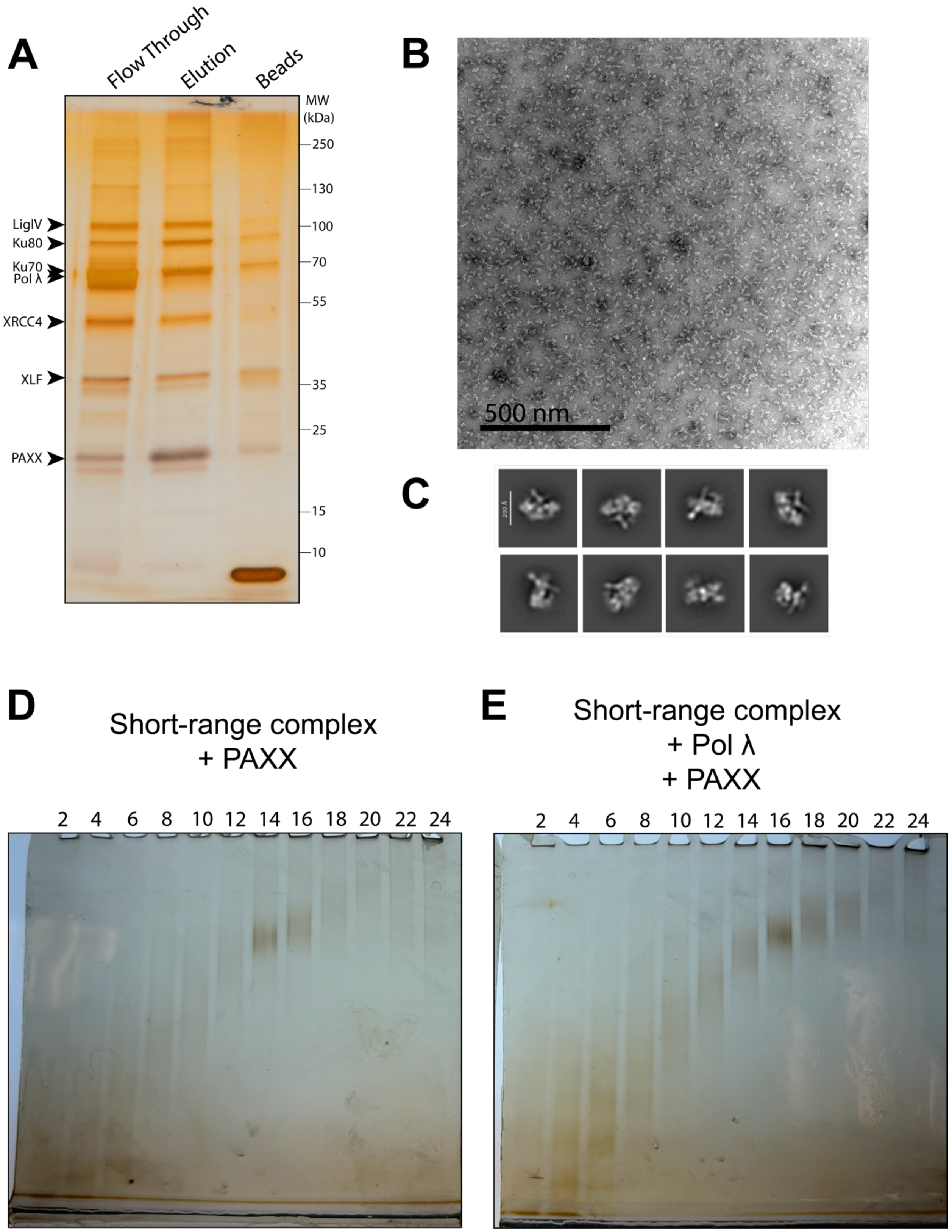
A
Flow Through
Elution
Beads
MW (kDa)
250
130
100
70
55
35
25
15
10
LigIV
Ku80
Ku70
Pol λ
XRCC4
XLF
PAXX
B
500 nm
C
D
Short-range complex
+ PAXX
2 4 6 8 10 12 14 16 18 20 22 24
E
Short-range complex
+ Pol λ
+ PAXX
2 4 6 8 10 12 14 16 18 20 22 24

**Supplemental Figure 1. Assembly of SR-Pol λ complex. (A)** Silver stain of a magnetic beads-based pulldown of the SR complex with Pol λ. **(B)** Negative stain EM of the SR-Pol λ complex. **(C)** 2D class averages of the SR-Pol λ complex from negative stain EM. **(D-E)** Non-denaturing PAGE analysis of GraFix sample preparations of the SR complex +/- Pol λ showing a clear peak shift from fractions 14-15 to 16-17.

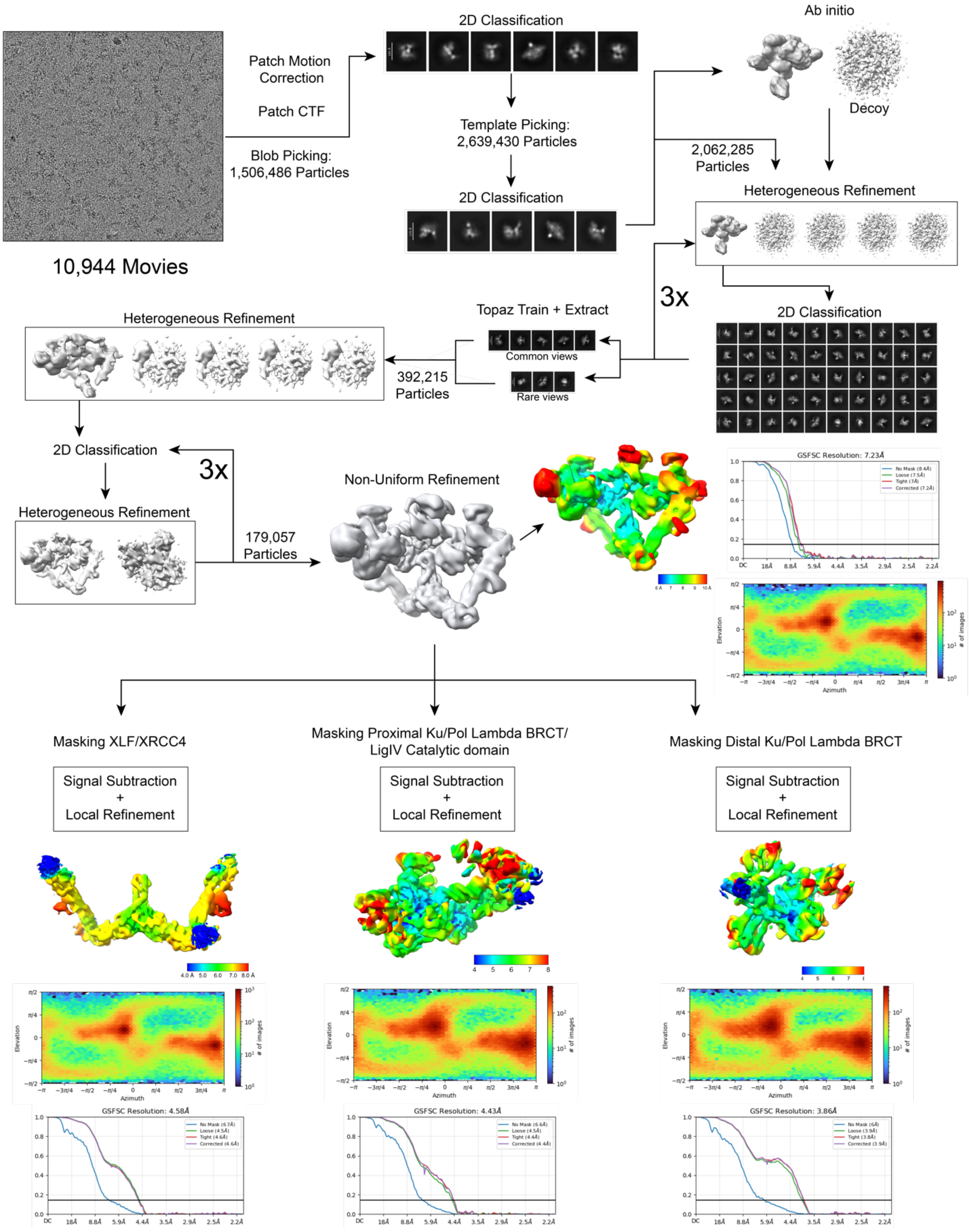

10,944 Movies
Patch Motion Correction
Patch CTF
Blob Picking: 1,506,486 Particles
2D Classification
Template Picking: 2,639,430 Particles
2D Classification
Ab initio
Decoy
2,062,285 Particles
Heterogeneous Refinement
3x
2D Classification
Topaz Train + Extract
Common views
Rare views
392,215 Particles
Heterogeneous Refinement
2D Classification
3x
Heterogeneous Refinement
179,057 Particles
Non-Uniform Refinement
GSFSC Resolution: 7.23Å
Masking XLF/XRCC4
Masking Proximal Ku/Pol Lambda BRCT/ LigIV Catalytic domain
Masking Distal Ku/Pol Lambda BRCT
Signal Subtraction + Local Refinement
Signal Subtraction + Local Refinement
Signal Subtraction + Local Refinement
GSFSC Resolution: 4.58Å
GSFSC Resolution: 4.43Å
GSFSC Resolution: 3.86Å

**Supplemental Figure 2. Data-processing scheme of the SR-Pol λ complex.** Flow chart of cryo-EM data processing. Gold-standard Fourier shell correlation (FSC) curves (cutoff 0.143) show the final calculated resolution of each individually refined body.

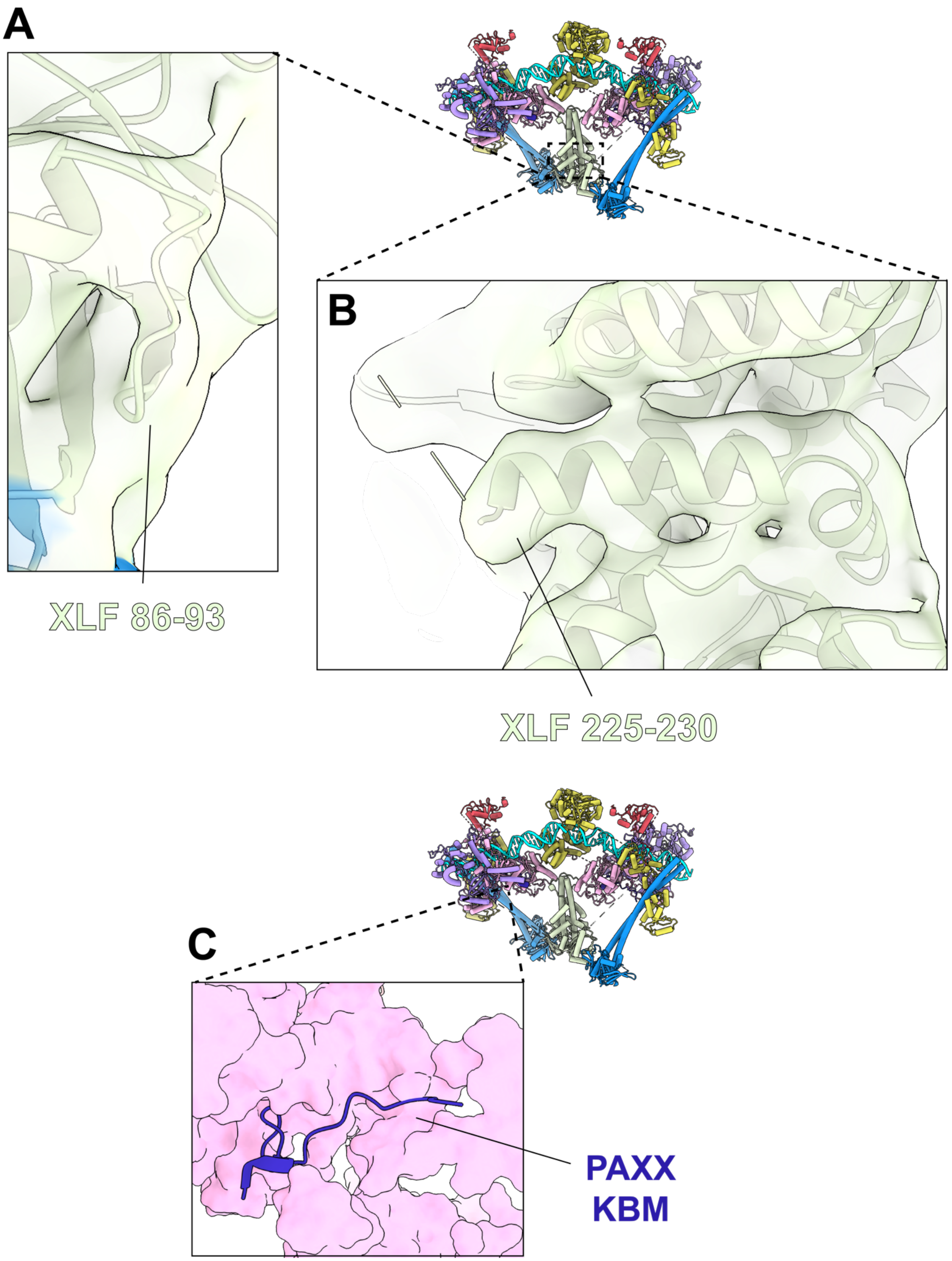
A
XLF 86-93
B
XLF 225-230
C
PAXX
KBM

**Supplemental Figure 3. Structural details of core NHEJ factors in SR-Pol λ complex. (A)** Magnified view of the newly modeled loop of XLF with corresponding residues noted. **(B)** Magnified view of newly modeled extended helix of XLF with corresponding residues noted. (**C)** Magnification of the PAXX-KBM interacting with Ku70.

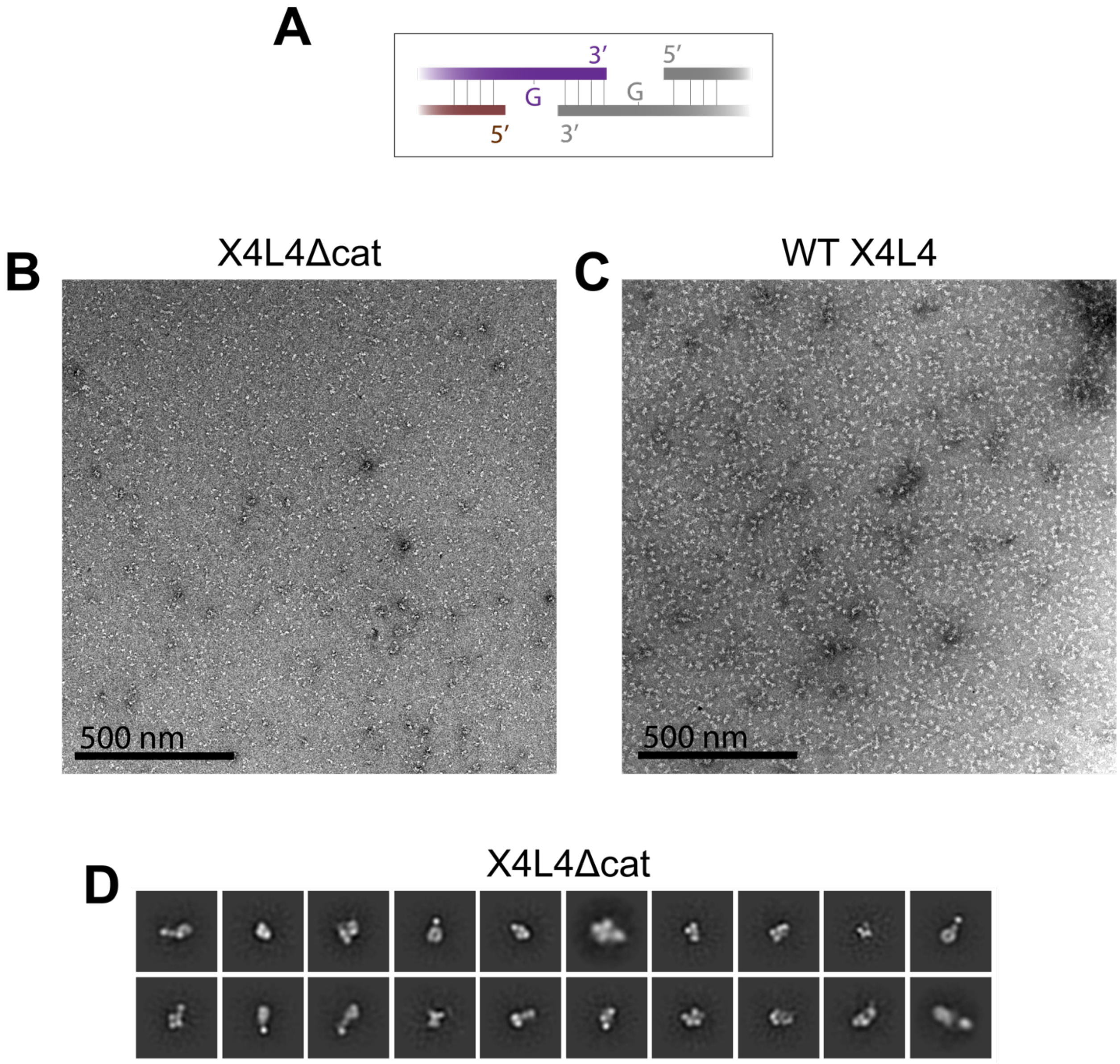

**Supplemental Figure 4. Negative stain of SR complex with X4L4Δcat. (A)** Diagram of ends of DNA substrate used – 4 nt complementarity flanked by 1 nt gaps. **(B)** Raw micrograph of SR complex assembled with X4L4Δcat. Particles are noticeably smaller than those observed in **(C) -** SR complex assembled with wild-type X4L4. **(D)** Representative 2D class averages for X4L4Δcat sample. No views of a fully assembled synaptic complex are observed.

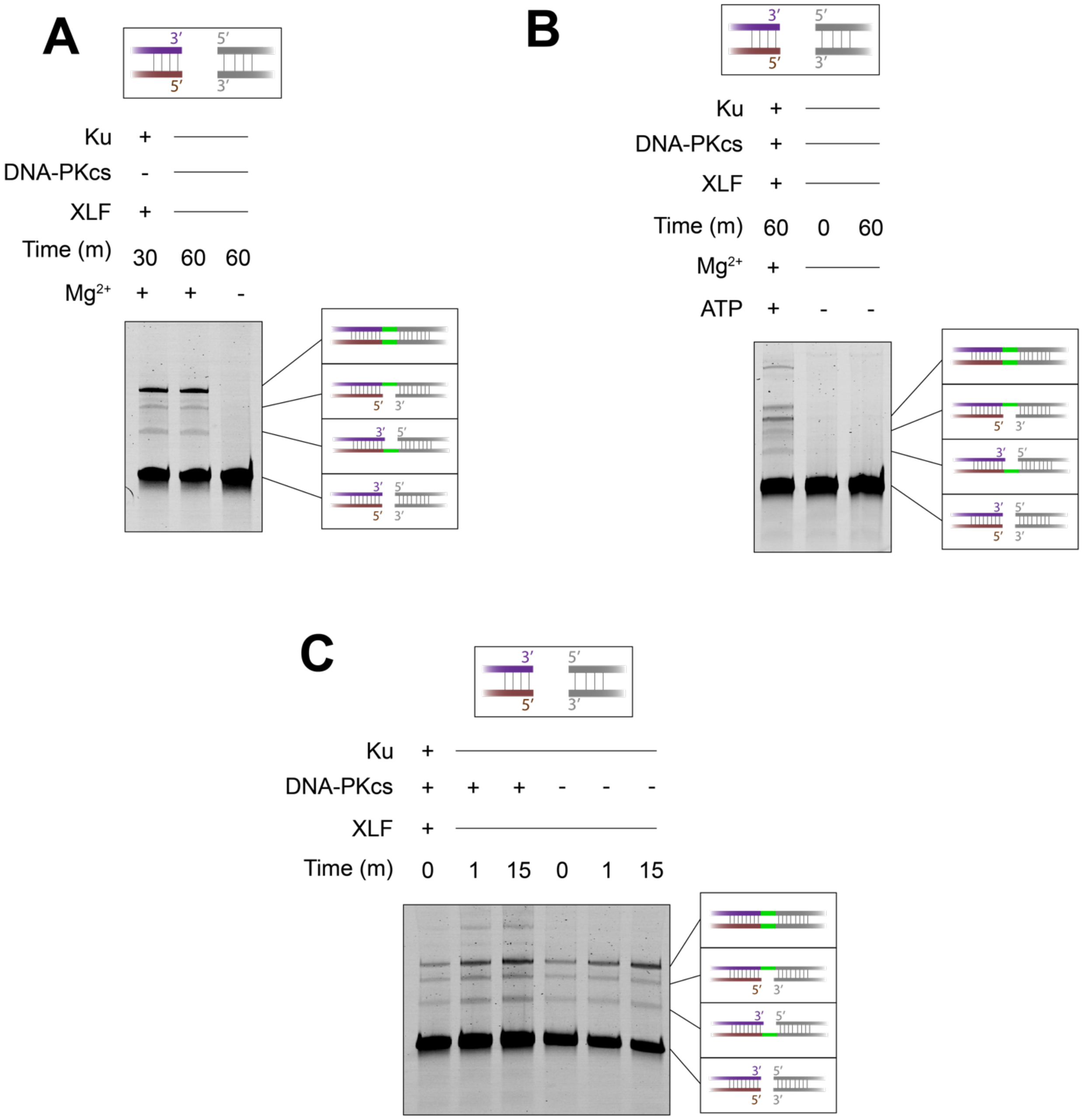

**Supplemental Figure 5. Validation of activity assay on directly ligatable substrates. (A)** Denaturing urea gel analysis of SR complex ligation activity demonstrating the attenuation of the reaction by addition/restriction of $Mg^{2+}$. **(B)** Assay of LR complex ligation. Compared to the SR complex, ligation is ATP dependent. **(C)** Time course of both LR and SR complex ligation reactions. Significant ligation is observed immediately after addition of $MgCl_2$ or ATP, indicating that repair can be carried out relatively quickly after the functional complex is assembled.

A

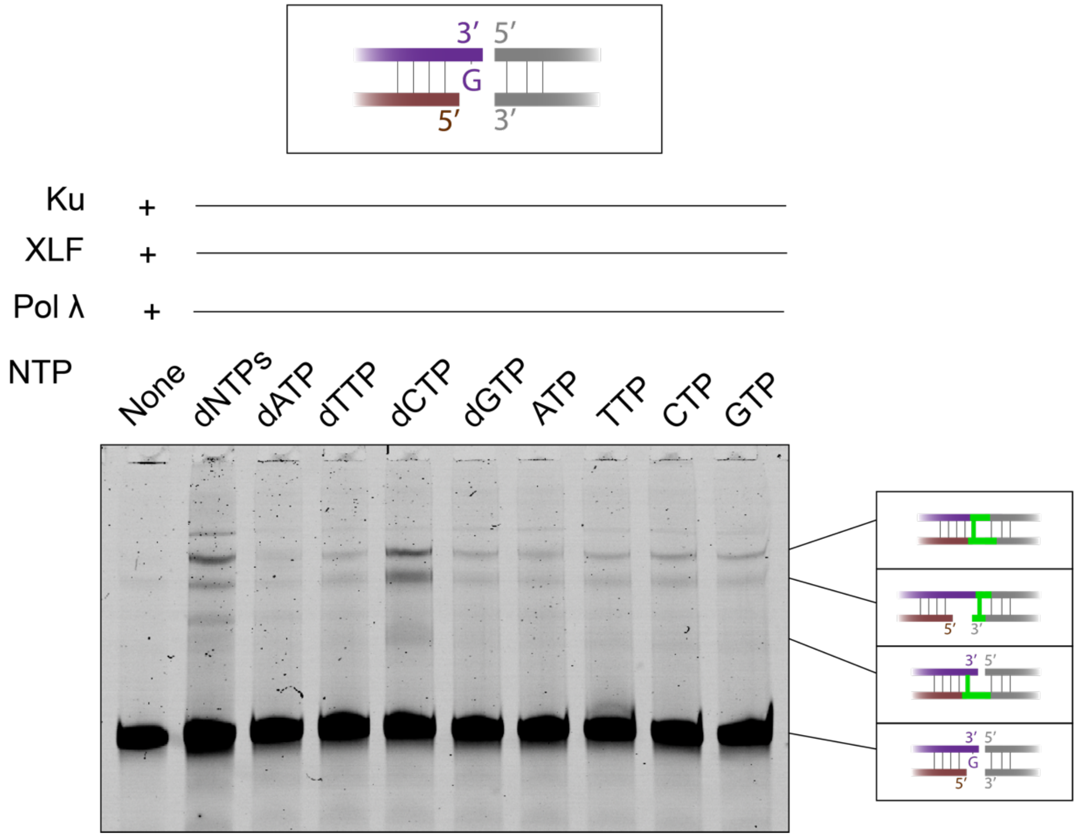

**Supplemental Figure 6. Comparison of NTP and dNTP usage by Pol λ. (A)** Comparison of Pol λ dependent ligation when supplied with different ribonucleotides and deoxyribonucleotides. dNTPs refers to a master mix of all dNTPs.

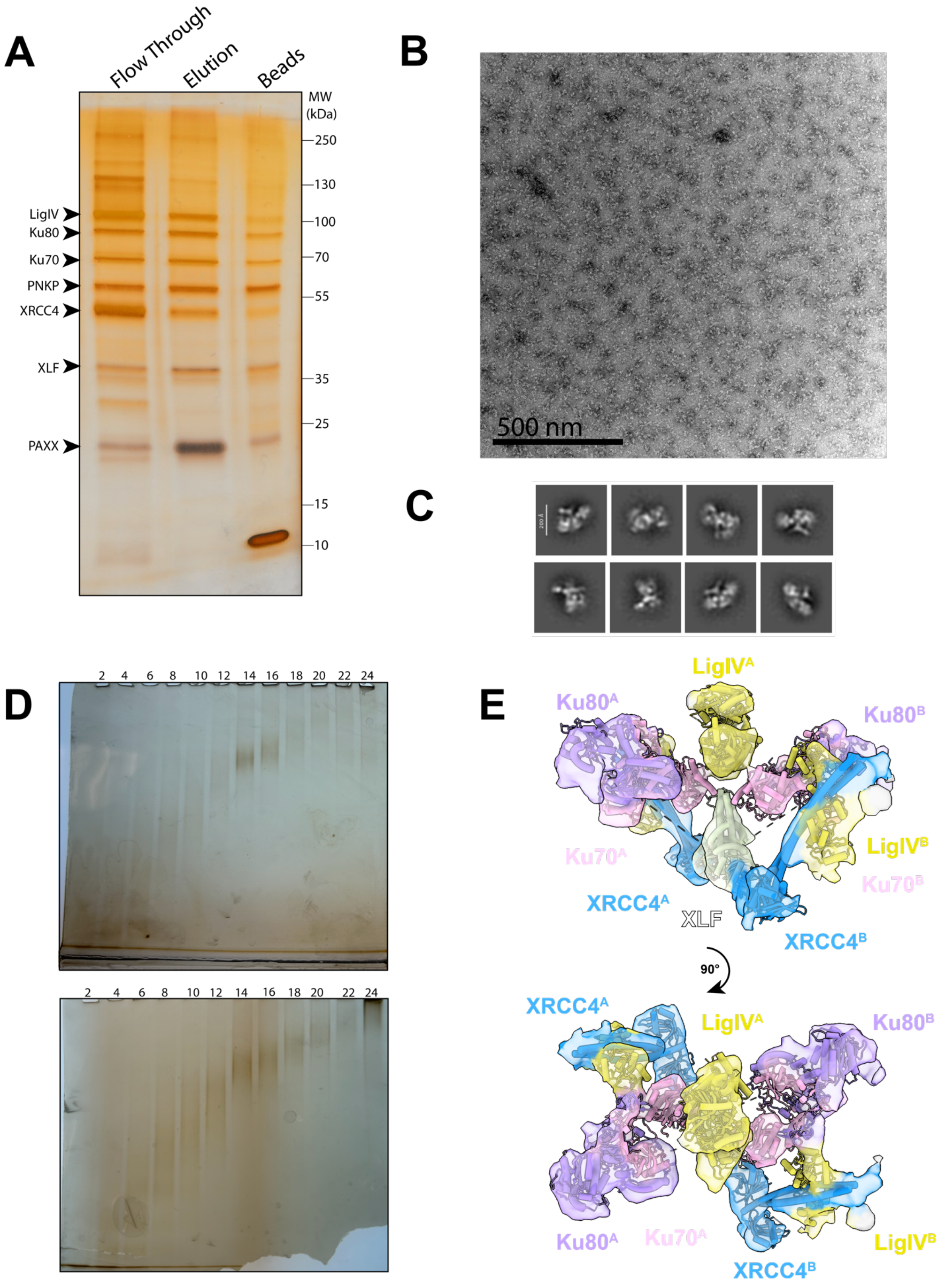
A
Flow Through
Elution
Beads
MW (kDa)
250
130
100
70
55
35
25
15
10
LigIV
Ku80
Ku70
PNKP
XRCC4
XLF
PAXX
B
500 nm
C
D
2 4 6 8 10 12 14 16 18 20 22 24
2 4 6 8 10 12 14 16 18 20 22 24
E
LigIV^A
Ku80^A
Ku80^B
Ku70^A
LigIV^B
Ku70^B
XRCC4^A
XLF
XRCC4^B
90°
XRCC4^A
LigIV^A
Ku80^B
Ku80^A
Ku70^A
XRCC4^B
LigIV^B

**Supplemental Figure 7. Assembly and negative stain EM of SR-PNKP complex.**

**(A)** Silver stain of a magnetic beads-based pulldown of the SR complex with PNKP. **(B)** Negative stain EM of the SR-PNKP complex. **(C)** 2D class averages of the SR-PNKP complex from negative stain EM. **(D)** Non-denaturing PAGE analysis of GraFix sample preparations of the SR complex +/- PNKP. **(E)** 3D reconstruction from negative stain EM of the SR-PNKP complex. No new density corresponding to PNKP is observed.

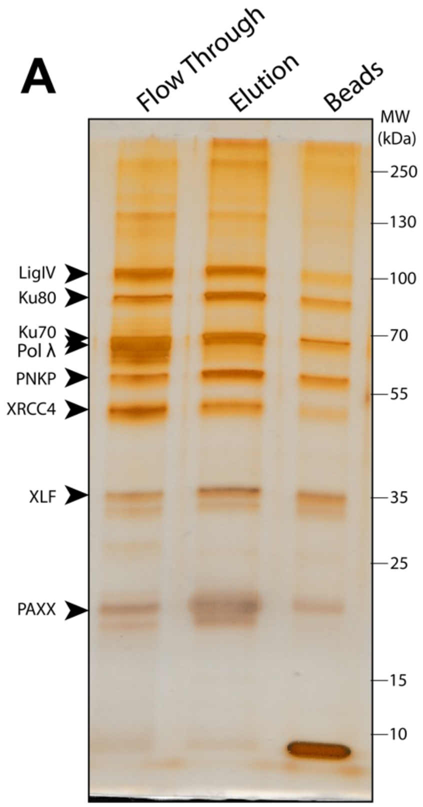

**Supplemental Figure 8. Pulldown of SR complex containing both Pol λ and PNKP.**

**(A)** Silver stain of a magnetic beads-based pulldown of the SR complex with Pol λ and PNKP.

**Table 1. List of oligonucleotides used in study**

| Construct | Fig(s) | Sran d | Sequence |
|---|---|---|---|
| Y45_3'ovh1 | 1,2 | A | ACCTCCCACTATTTTTCCGGGCAAGCTCGATCCCCGAGCTTCTTTGA ACTTTGTTGTCTGGAGTCTCTGTGGCAATCGATGATGTG |
| | | B | /5phos/ACATCATCGATTGCCACAGAGACTCCAGACAACAAAGTTC AAAGATGCCCGG |
| | | RNA bait | UAGUGGGAGGU/3BioTEG/ |
| Y30_blunt_ end | 3B, 4B-D, Sup. 7A-C | A | ACCTCCCACTATTTTTCCGGGCAAGCTCGATCCCCGAGCTTCTAAG AACTCTGATGTCAGTAGATTACAC |
| | | B | /5phos/GTGTAATCTACTGACATCAGAGTTCTTAGATGCCCGG |
| | | C | /5phos/ACGTAGACTGCTGACATCAGAGTTCTTAGATGCCCGGTTT /iCy5/TTCCGGGCAAGCTCGATCCCCGAGCTTCTAAGAACTCTGAT GTCAGCAGTCTACGT |
| | | RNA bait | UAGUGGGAGGU/3BioTEG/ |
| Y30_C4 | 3E-F, 4C, 4E, 5D | A | ACCTCCCACTATTTTTCCGGGCAAGCTCGATCCCCGAGCTTCTAAG AACTCTGATGTCAGCAGTCTACGTGTTC |
| | | B | /5phos/CGTAGACTGCTGACATCAGAGTTCTTAGATGCCCGG |
| | | C | /5phos/ACGTAGACTGCTGACATCAGAGTTCTTAGATGCCCGGTTT /iCy5/TTCCGGGCAAGCTCGATCCCCGAGCTTCTAAGAACTCTGAT GTCAGCAGTCTACGTGAAC |
| | | RNA bait | UAGUGGGAGGU/3BioTEG/ |
| Y30_3'ovh1 | 3C | A | ACCTCCCACTATTTTTCCGGGCAAGCTCGATCCCCGAGCTTCTAAG AACTCTGATGTCAGCAGTCTACGTGT |
| | | B | /5phos/CACGTAGACTGCTGACATCAGAGTTCTTAGATGCCCGG |
| | | C | /5phos/ACGTAGACTGCTGACATCAGAGTTCTTAGATGCCCGGTTT /iCy5/TTCCGGGCAAGCTCGATCCCCGAGCTTCTAAGAACTCTGAT GTCAGCAGTCTACGT |
| | | RNA bait | UAGUGGGAGGU/3BioTEG/ |

| | | | |
|---|---|---|---|
| Y30_3'ovh2 | 3C | A | ACCTCCCACTATTTTTCCGGGCAAGCTCGATCCCCGAGCTTCTAAGAACTCTGATGTCAGCAGTCTACGTGT |
| | | B | /5phos/ACGTAGACTGCTGACATCAGAGTTCTTAGATGCCCGG |
| | | C | /5phos/ACGTAGACTGCTGACATCAGAGTTCTTAGATGCCCGGTTT/iCy5/TTCCGGGCAAGCTCGATCCCCGAGCTTCTAAGAACTCTGATGTCAGCAGTCTACGT |
| | | RNA bait | UAGUGGGAGGU/3BioTEG/ |
| Y30_3'ovh3 | 3C | A | ACCTCCCACTATTTTTCCGGGCAAGCTCGATCCCCGAGCTTCTAAGAACTCTGATGTCAGCAGTCTACGTGTT |
| | | B | /5phos/ACGTAGACTGCTGACATCAGAGTTCTTAGATGCCCGG |
| | | C | /5phos/ACGTAGACTGCTGACATCAGAGTTCTTAGATGCCCGGTTT/iCy5/TTCCGGGCAAGCTCGATCCCCGAGCTTCTAAGAACTCTGATGTCAGCAGTCTACGT |
| | | RNA bait | UAGUGGGAGGU/3BioTEG/ |
| Y30_3'ovh4 | 3C | A | ACCTCCCACTATTTTTCCGGGCAAGCTCGATCCCCGAGCTTCTAAGAACTCTGATGTCAGCAGTCTACGTGTTC |
| | | B | /5phos/ACGTAGACTGCTGACATCAGAGTTCTTAGATGCCCGG |
| | | C | /5phos/ACGTAGACTGCTGACATCAGAGTTCTTAGATGCCCGGTTT/iCy5/TTCCGGGCAAGCTCGATCCCCGAGCTTCTAAGAACTCTGATGTCAGCAGTCTACGT |
| | | RNA bait | UAGUGGGAGGU/3BioTEG/ |
| Y30_5'ovh1 | 3D | A | ACCTCCCACTATTTTTCCGGGCAAGCTCGATCCCCGAGCTTCTAAGAACTCTGATGTCAGCAGTCTACGTGT |
| | | B | /5phos/GACACGTAGACTGCTGACATCAGAGTTCTTAGATGCCCGG |
| | | C | /5phos/ACGTAGACTGCTGACATCAGAGTTCTTAGATGCCCGGTTT/iCy5/TTCCGGGCAAGCTCGATCCCCGAGCTTCTAAGAACTCTGATGTCAGCAGTCTACGT |
| | | RNA bait | UAGUGGGAGGU/3BioTEG/ |
| Y30_5'ovh2 | 3D | A | ACCTCCCACTATTTTTCCGGGCAAGCTCGATCCCCGAGCTTCTAAGAACTCTGATGTCAGCAGTCTACGTGT |

| | | | |
|---|---|---|---|
| | | B | /5phos/CGACACGTAGACTGCTGACATCAGAGTTCTTAGATGCCCGG |
| | | C | /5phos/ACGTAGACTGCTGACATCAGAGTTCTTAGATGCCCGGTTT/iCy5/TTCCGGGCAAGCTCGATCCCCGAGCTTCTAAGAACTCTGATGTCAGCAGTCTACGT |
| | | RNA bait | UAGUGGGAGGU/3BioTEG/ |
| Y30_5'ovh3 | 3D | A | ACCTCCCACTATTTTTCCGGGCAAGCTCGATCCCCGAGCTTCTAAGAACTCTGATGTCAGCAGTCTACGTG |
| | | B | /5phos/GAACACGTAGACTGCTGACATCAGAGTTCTTAGATGCCCGG |
| | | C | /5phos/ACGTAGACTGCTGACATCAGAGTTCTTAGATGCCCGGTTT/iCy5/TTCCGGGCAAGCTCGATCCCCGAGCTTCTAAGAACTCTGATGTCAGCAGTCTACGT |
| | | RNA bait | UAGUGGGAGGU/3BioTEG/ |
| Y30_5'ovh4 | 3D | A | ACCTCCCACTATTTTTCCGGGCAAGCTCGATCCCCGAGCTTCTAAGAACTCTGATGTCAGCAGTCTACGT |
| | | B | /5phos/GAACACGTAGACTGCTGACATCAGAGTTCTTAGATGCCCGG |
| | | C | /5phos/ACGTAGACTGCTGACATCAGAGTTCTTAGATGCCCGGTTT/iCy5/TTCCGGGCAAGCTCGATCCCCGAGCTTCTAAGAACTCTGATGTCAGCAGTCTACGT |
| | | RNA bait | UAGUGGGAGGU/3BioTEG/ |
| Y30_C3_doublegap | 3E-F | A | ACCTCCCACTATTTTTCCGGGCAAGCTCGATCCCCGAGCTTCTAAGAACTCTGATGTCAGCAGTCTACGTGTT |
| | | B | /5phos/CGTAGACTGCTGACATCAGAGTTCTTAGATGCCCGG |
| | | C | /5phos/ACGTAGACTGCTGACATCAGAGTTCTTAGATGCCCGGTTT/iCy5/TTCCGGGCAAGCTCGATCCCCGAGCTTCTAAGAACTCTGATGTCAGCAGTCTACGTGAAC |
| | | RNA bait | UAGUGGGAGGU/3BioTEG/ |
| Y30_bluntend_3'Pho/5'OH | 3B, 3D | A | ACCTCCCACTATTTTTCCGGGCAAGCTCGATCCCCGAGCTTCTAAGAACTCTGATGTCAGCAGTCTACGT/3Phos/ |
| | | B | ACGTAGACTGCTGACATCAGAGTTCTTAGATGCCCGG |

| | | | |
|---|---|---|---|
| | | C | /5phos/ACGTAGACTGCTGACATCAGAGTTCTTAGATGCCCGGTTT/iCy5/TTCCGGGCAAGCTCGATCCCCGAGCTTCTAAGAACTCTGATGTCAGCAGTCTACGT |
| | | RNA bait | UAGUGGGAGGU/3BioTEG/ |
| Y30_C4_5'OH | 3C, 3E | A | ACCTCCCACTATTTTTCCGGGCAAGCTCGATCCCCGAGCTTCTAAGAACTCTGATGTCAGCAGTCTACGTGTTC |
| | | B | ACGTAGACTGCTGACATCAGAGTTCTTAGATGCCCGG |
| | | C | /5phos/ACGTAGACTGCTGACATCAGAGTTCTTAGATGCCCGGTTT/iCy5/TTCCGGGCAAGCTCGATCCCCGAGCTTCTAAGAACTCTGATGTCAGCAGTCTACGTGAAC |
| | | RNA bait | UAGUGGGAGGU/3BioTEG/ |
| Y30_bluntend_5'OH | 3C | A | ACCTCCCACTATTTTTCCGGGCAAGCTCGATCCCCGAGCTTCTAAGAACTCTGATGTCAGCAGTCTACGT |
| | | B | ACGTAGACTGCTGACATCAGAGTTCTTAGATGCCCGG |
| | | C | /5phos/ACGTAGACTGCTGACATCAGAGTTCTTAGATGCCCGGTTT/iCy5/TTCCGGGCAAGCTCGATCCCCGAGCTTCTAAGAACTCTGATGTCAGCAGTCTACGT |
| | | RNA bait | UAGUGGGAGGU/3BioTEG/ |
| Y30_bluntend_3'Pho | 3D | A | ACCTCCCACTATTTTTCCGGGCAAGCTCGATCCCCGAGCTTCTAAGAACTCTGATGTCAGCAGTCTACGT/3Phos/ |
| | | B | /5phos/ACGTAGACTGCTGACATCAGAGTTCTTAGATGCCCGG |
| | | C | /5phos/ACGTAGACTGCTGACATCAGAGTTCTTAGATGCCCGGTTT/iCy5/TTCCGGGCAAGCTCGATCCCCGAGCTTCTAAGAACTCTGATGTCAGCAGTCTACGT |
| | | RNA bait | UAGUGGGAGGU/3BioTEG/ |
| Y30_C4_3'Pho | 3E | A | ACCTCCCACTATTTTTCCGGGCAAGCTCGATCCCCGAGCTTCTAAGAACTCTGATGTCAGCAGTCTACGTGTTC/3'Pho/ |
| | | B | /5phos/ACGTAGACTGCTGACATCAGAGTTCTTAGATGCCCGG |
| | | C | /5phos/ACGTAGACTGCTGACATCAGAGTTCTTAGATGCCCGGTTT/iCy5/TTCCGGGCAAGCTCGATCCCCGAGCTTCTAAGAACTCTGATGTCAGCAGTCTACGTGAAC |

| | | | |
|---|---|---|---|
| | | RNA bait | UAGUGGGAGGU/3BioTEG/ |
| Y30_C4_3'Pho/5'OH | 3E | A | ACCTCCCACTATTTTTCCGGGCAAGCTCGATCCCCGAGCTTCTAAGAACTCTGATGTCAGCAGTCTACGTGTTC/3’Pho/ |
| | | B | ACGTAGACTGCTGACATCAGAGTTCTTAGATGCCCGG |
| | | C | /5phos/ACGTAGACTGCTGACATCAGAGTTCTTAGATGCCCGGTTT/iCy5/TTCCGGGCAAGCTCGATCCCCGAGCTTCTAAGAACTCTGATGTCAGCAGTCTACGTGAAC |
| | | RNA bait | UAGUGGGAGGU/3BioTEG/ |
| Y30_5'OH_3'ovh4 | 5A | A | ACCTCCCACTATTTTTCCGGGCAAGCTCGATCCCCGAGCTTCTAAGAACTCTGATGTCAGCAGTCTACGT |
| | | B | ACGTAGACTGCTGACATCAGAGTTCTTAGATGCCCGG |
| | | C | /5phos/ACGTAGACTGCTGACATCAGAGTTCTTAGATGCCCGGTTT/iCy5/TTCCGGGCAAGCTCGATCCCCGAGCTTCTAAGAACTCTGATGTCAGCAGTCTACGT |
| | | RNA bait | UAGUGGGAGGU/3BioTEG/ |
| Y30_3'Pho_3'ovh4 | 5B | A | ACCTCCCACTATTTTTCCGGGCAAGCTCGATCCCCGAGCTTCTAAGAACTCTGATGTCAGCAGTCTACGT/3Phos/ |
| | | B | /5phos/ACGTAGACTGCTGACATCAGAGTTCTTAGATGCCCGG |
| | | C | /5phos/ACGTAGACTGCTGACATCAGAGTTCTTAGATGCCCGGTTT/iCy5/TTCCGGGCAAGCTCGATCCCCGAGCTTCTAAGAACTCTGATGTCAGCAGTCTACGT |
| | | RNA bait | UAGUGGGAGGU/3BioTEG/ |
| Y30_3'Pho/5'OH_3'ovh4 | 5C | A | ACCTCCCACTATTTTTCCGGGCAAGCTCGATCCCCGAGCTTCTAAGAACTCTGATGTCAGCAGTCTACGTGTTC/3’Pho/ |
| | | B | ACGTAGACTGCTGACATCAGAGTTCTTAGATGCCCGG |
| | | C | /5phos/ACGTAGACTGCTGACATCAGAGTTCTTAGATGCCCGGTTT/iCy5/TTCCGGGCAAGCTCGATCCCCGAGCTTCTAAGAACTCTGATGTCAGCAGTCTACGT |
| | | RNA bait | UAGUGGGAGGU/3BioTEG/ |

| | | | |
|---|---|---|---|
| Y30_C3_gap_5'OH | 5D | A | ACCTCCCACTATTTTTCCGGGCAAGCTCGATCCCCGAGCTTCTAAGAACTCTGATGTCAGCAGTCTACGTGTT |
| | | B | ACGTAGACTGCTGACATCAGAGTTCTTAGATGCCCGG |
| | | C | /5phos/ACGTAGACTGCTGACATCAGAGTTCTTAGATGCCCGGTTT/iCy5/TTCCGGGCAAGCTCGATCCCCGAGCTTCTAAGAACTCTGATGTCAGCAGTCTACGTGAAC |
| | | RNA bait | UAGUGGGAGGU/3BioTEG/ |
| Y30_C3_gap_3'Pho | 5E | A | ACCTCCCACTATTTTTCCGGGCAAGCTCGATCCCCGAGCTTCTAAGAACTCTGATGTCAGCAGTCTACGTGTTC/3'Pho/ |
| | | B | /5phos/CGTAGACTGCTGACATCAGAGTTCTTAGATGCCCGG |
| | | C | /5phos/ACGTAGACTGCTGACATCAGAGTTCTTAGATGCCCGGTTT/iCy5/TTCCGGGCAAGCTCGATCCCCGAGCTTCTAAGAACTCTGATGTCAGCAGTCTACGTGAAC |
| | | RNA bait | UAGUGGGAGGU/3BioTEG/ |
| Y45_C4_gap | Sup. 4 | A (left) | ACCTCCCACTATTTTTCCGGGCAAGCTCGATCCCCGAGCTTCTTTGAACTTTGTTGTCTGGAGTCTCTGTGGCAATCGATGATGTGGTTC |
| | | B (left) | /5phos/ACATCATCGATTGCCACAGAGACTCCAGACAACAAAGTTCAAAGATGCCCGG |
| | | A (right) | CCGGGCAAGCTCGATCCCCGAGCTTCTTTGAACTTTGTTGTCTGGAGTCTCTGTGGCAATCGATGATGTGGAAC |
| | | B (right) | /5phos/ACATCATCGATTGCCACAGAGACTCCAGACAACAAAGTTCAAAGATGCCCGG |
| | | RNA bait | UAGUGGGAGGU/3BioTEG/ |

### Table 2. Cryo-EM data collection, refinement and validation statistics

| | #1 Short-range Pol λ complext (EMDB-xxxx) (PDB xxxx) |
|---|---|
| **Data collection and processing** | |
| Magnification | 30k |
| Voltage (kV) | 300 |
| Electron exposure (e–/Å$^2$) | |
| Defocus range (μm) | -1.5 to -4.0 |
| Pixel size (Å) | 1.056 |
| Symmetry imposed | C1 |
| Initial particle images (no.) | 1,506,486 |
| Final particle images (no.) | 179,057 |
| Map resolution (Å) FSC threshold = 0.142 | 7.23 (overall), 4.58 (XRCC4-LigIV, XLF scaffold), 4.43 (proximal Ku, Pol λ BRCT, LigIV catalytic domain), 3.86 (distal Ku, Pol λ BRCT) |
| Map resolution range (Å) | 3.86 to >10 |
| **Refinement** | |
| Initial model used (PDB code) | 7LSY |
| Model resolution (Å) FSC threshold | 4.58 (XRCC4-LigIV, XLF scaffold), 4.43 (proximal Ku, Pol λ BRCT, LigIV catalytic domain), 3.86 (distal Ku, Pol λ BRCT) 0.143 |
| Model resolution range (Å) | n/a |
| Map sharpening *B* factor (Å$^2$) | -615 (overall), -169 (XRCC4-LigIV, XLF scaffold), -147 (proximal Ku, Pol λ BRCT, LigIV catalytic domain), -123 (distal Ku, Pol λ BRCT) |
| Model composition | |
| Non-hydrogen atoms | 77,946 |
| Protein residues | 4,632 (Nucleotide: 136) |
| Ligands | 0 |
| *B* factors (Å$^2$) | Not estimated |
| Protein | |
| Ligand | |
| R.m.s. deviations | |
| Bond lengths (Å) | 0.013 |
| Bond angles (°) | 1.614 |
| Validation | |
| MolProbity score | 2.20 |
| Clashscore | 13.25 |
| Poor rotamers (%) | 1.42 |
| Ramachandran plot | |
| Favored (%) | 92.96 |
| Allowed (%) | 7.04 |
| Disallowed (%) | 0.00 |
